# Reproducibility of functional brain alterations in major depressive disorder: evidence from a multisite resting-state functional MRI study with 1,434 individuals

**DOI:** 10.1101/524496

**Authors:** Mingrui Xia, Tianmei Si, Xiaoyi Sun, Qing Ma, Bangshan Liu, Li Wang, Jie Meng, Miao Chang, Xiaoqi Huang, Ziqi Chen, Yanqing Tang, Ke Xu, Qiyong Gong, Fei Wang, Jiang Qiu, Peng Xie, Lingjiang Li, Yong He, DIDA-Major Depressive Disorder Working Group

**Author notes:** These authors contributed equally to this work (Mingrui Xia, Tianmei Si). These authors contributed equally to this work (Qiyong Gong, Fei Wang, Jiang Qiu, Peng Xie, Lingjiang Li, Yong He). Correspondence: Yong He, PhD., State Key Laboratory of Cognitive Neuroscience and Learning, Beijing Key Laboratory of Brain Imaging and Connectomics, IDG/McGovern Institute for Brain Research, Beijing Normal University, Beijing, 100875, China. Or Lingjiang Li, MD, PhD, Mental Health Institute, Second Xiangya Hospital of Central South University, Changsha, 410011, China.

## Abstract

Resting-state functional MRI (R-fMRI) studies have demonstrated widespread alterations in brain function in patients with major depressive disorder (MDD). However, a clear and consistent conclusion regarding a repeatable pattern of MDD-relevant alterations is still limited due to the scarcity of large-sample, multisite datasets. Here, we address this issue by including a large R-fMRI dataset with 1,434 participants (709 patients with MDD and 725 healthy controls) from five centers in China. Individual functional activity maps that represent very local to long-range connections are computed using the amplitude of low-frequency fluctuations, regional homogeneity and distance-related functional connectivity strength. The reproducibility analyses involve different statistical strategies, global signal regression, across-center consistency, clinical variables, and sample size. We observed significant hypoactivity in the orbitofrontal, sensorimotor, and visual cortices and hyperactivity in the frontoparietal cortices in MDD patients compared to the controls. These alterations are not affected by different statistical analysis strategies, global signal regression and medication status and are generally reproducible across centers. However, these between-group differences are partially influenced by the episode status and the age of disease onset in patients, and the brain-clinical variable relationship exhibits poor cross-center reproducibility. Bootstrap analyses reveal that at least 400 subjects in each group are required to replicate significant alterations (an extent threshold of *P*<.05 and a height threshold of *P*<.001) at 50% reproducibility. Together, these results highlight reproducible patterns of functional alterations in MDD and relevant influencing factors, which provides crucial guidance for future neuroimaging studies of this disorder.

## Introduction

Major depressive disorder (MDD) is the leading contributor to years-of-life lived with disability, and it is characterized by mood disturbances, loss of interest in activities and deficits in cognitive functions, resulting in increasing economic and social burdens (Kessler et al., 2007; Murray et al., 2012). Many previous studies of MDD have revealed structural and functional alterations in the brain that have substantially enhanced our understanding of the neurobiological substrates underlying the behavioral deficits in patients with MDD. However, the pathophysiological mechanism of MDD is incompletely understood (DeRubeis et al., 2008; Gong and He, 2015).

Over the past two decades, rapid advances in resting-state functional MRI (R-fMRI) have provided an unprecedented opportunity for the noninvasive investigation of functional architecture in spontaneous or intrinsic brain activities within and between regions (Biswal et al., 1995; Fox and Raichle, 2007; Wang et al., 2015a). Relating to depression, many R-fMRI studies have documented widespread functional alterations in patients with MDD, involving the primary sensorimotor (Kuhn and Gallinat, 2013; Wang et al., 2014a; Zhang et al., 2011) and visual (Kaiser et al., 2015; Kuhn and Gallinat, 2013) cortices, medial/lateral prefrontal (Greicius et al., 2007; Kaiser et al., 2015; Kuhn and Gallinat, 2013; Lui et al., 2011; Sheline et al., 2010; Wang et al., 2014a; Wang et al., 2015b; Zhang et al., 2011; Zhu et al., 2012) and parietal (Kaiser et al., 2015; Sheline et al., 2010; Wang et al., 2014a; Zhang et al., 2011; Zhu et al., 2012) cortices, and subcortical areas (Anand et al., 2009; Greicius et al., 2007; Kaiser et al., 2015; Kuhn and Gallinat, 2013; Lui et al., 2011; Wang et al., 2014a; Wang et al., 2015b); these areas cover approximately the whole brain. It is important to note that a clear and consistent conclusion regarding whether these functional alterations are reliable and can be reproduced in patients with MDD is still limited. The causes of this phenomenon might result from the limited statistical power of a small research sample, different patient recruitment criteria (e.g., cultural background and diagnostic criteria), different imaging protocols (e.g., MRI scanners and imaging parameters) and different analysis strategies (e.g., preprocessing procedures and functional brain measures) across studies (Button et al., 2013; Gong and He, 2015).

With the continuous emergence of datasets or consortiums with large sample sizes, high-quality data and multidimensional variables, functional imaging research of the brain is entering the era of “big data” (Poldrack and Gorgolewski, 2014; Xia and He, 2017). These data are extremely important for identifying reliable patterns of functional brain alterations in psychiatric disorders such as MDD due to the benefits of greater statistical power and across-center validations. For instance, using a large-sample R-fMRI dataset of 421 patients with MDD and 488 healthy controls (HCs) from three research sites, Cheng *et al*. reported abnormal functional connectivity in the medial reward and lateral nonreward circuits of the orbitofrontal cortex in patients with MDD (Cheng et al., 2016). Using R-fMRI data from 458 patients with MDD and 730 HCs from two sites, Drysdale *et al*. divided depressed patients into different neurophysiological subtypes according to their connectivity dysfunctions and successfully predicted their differential responses to transcranial magnetic stimulation therapy (Drysdale et al., 2017). However, large-sample, multicenter neuroimaging studies aiming to evaluate the reproducibility of functional alterations in patients with MDD are still lacking.

Here, we collected a large R-fMRI dataset of 1,434 participants, including 709 patients with MDD and 725 HCs, from five centers in China, thus ensuring homogeneity in genetic and cultural backgrounds in the same race. We calculated individual functional activity maps that represent very local to long-range connections using three frequently used R-fMRI measures, including the amplitude of low-frequency fluctuations (ALFF) (Zang et al., 2007), regional homogeneity (ReHo) (Zang et al., 2004) and distance-dependent functional connectivity strength (FCS) (Buckner et al., 2009; Dai et al., 2015; Liang et al., 2013; Liao et al., 2013; Xia et al., 2018). We then examined functional brain alterations in patients with MDD, followed by reproducibility analyses involving different preprocessing and statistical analysis strategies with cross-center validation.

## Materials and Methods

### Participants

This study included 1,558 participants (782 patients with MDD and 776 HCs) who were recruited from five research centers in China through the Disease Imaging Data Archiving - Major Depressive Disorder Working Group (DIDA-MDD). All patients were diagnosed according to the Diagnostic and Statistical Manual of Mental Disorders IV (DSM-IV) criteria for MDD (First et al., 1997). The severity of depression was rated using the Hamilton Depression Rating Scale (HDRS) (Williams, 1988). Quality control was performed for both clinical and imaging data, including the presence of demographic information, completeness of R-fMRI scan, inconsistency in key scan parameters, errors in reading raw Digital Imaging and Communications in Medicine (DICOM) data, abnormalities in anatomical brain images, head motion, and coverage of the whole brain. The final sample included 1,434 participants (709 patients with MDD and 725 HCs). For detailed inclusion and exclusion criteria of data quality control in each center, see Supplementary Information. The study was approved by the ethics committees of each center, and written informed consent was obtained from each participant. Table 1 illustrates the demographics, clinical characteristics, and imaging data quality.

**Table 1.**
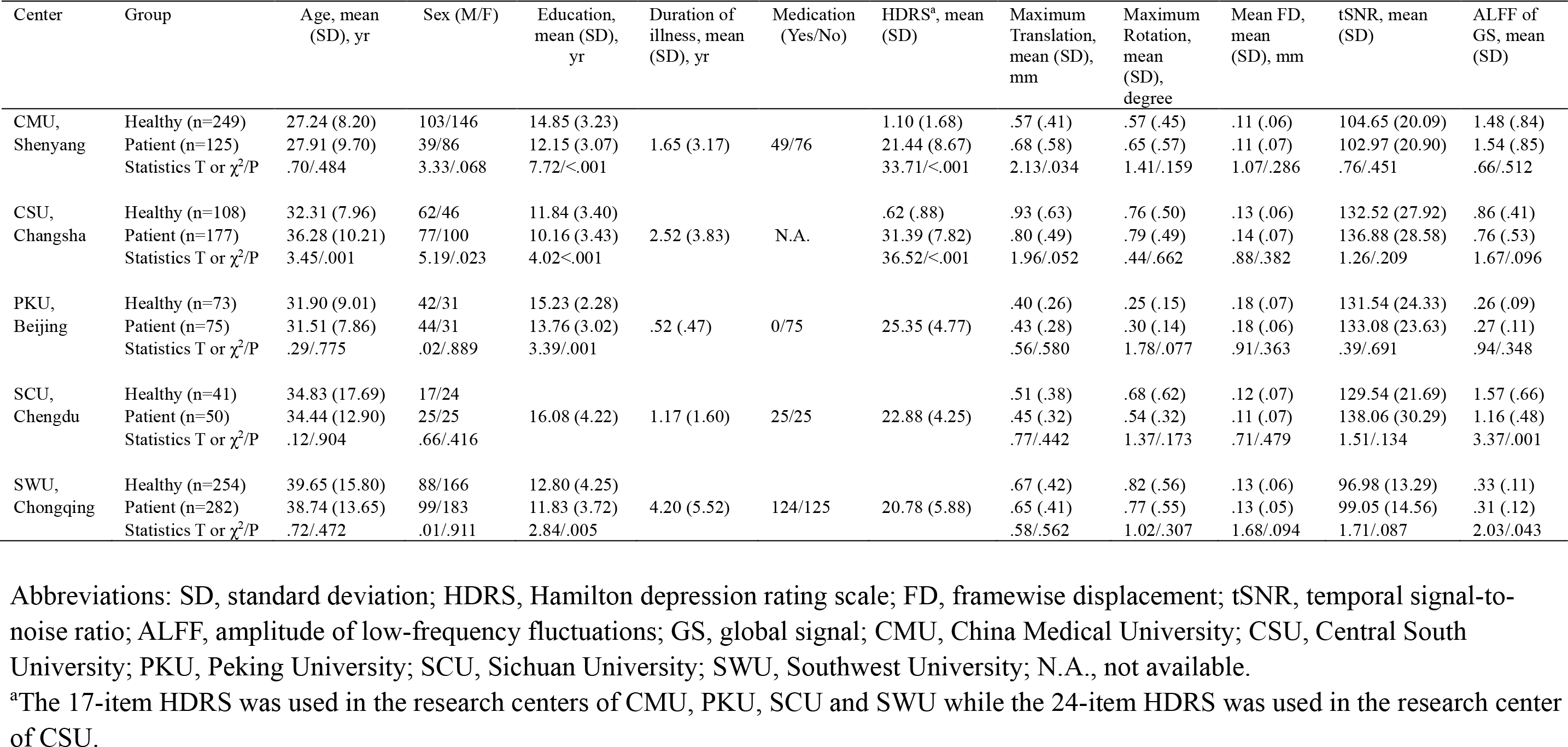
Demographic, clinical and imaging quality characteristics

### Image Acquisition

All R-fMRI data were obtained on 3.0-T MRI scanners with gradient-echo planar imaging sequences. During the scan, the participants were instructed to keep their eyes closed without falling asleep and move as little as possible. Detailed scanning parameters for each center are listed below (see Table 2 for summary).

**Table 2.**
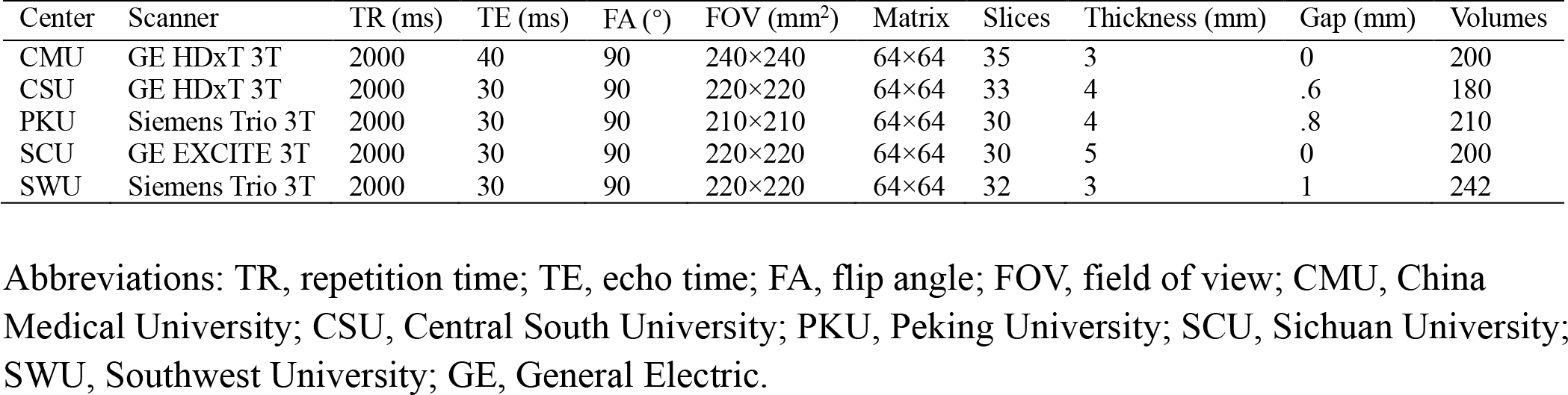
Scan parameters of R-fMRI data in each center

| Center | Scanner | TR (ms) | TE (ms) | FA (°) | FOV (mm<sup>2</sup>) | Matrix | Slices | Thickness (mm) | Gap (mm) | Volumes |
| --- | --- | --- | --- | --- | --- | --- | --- | --- | --- | --- |
| CMU | GE HDxT 3T | 2000 | 40 | 90 | 240×240 | 64×64 | 35 | 3 | 0 | 200 |
| CSU | GE HDxT 3T | 2000 | 30 | 90 | 220×220 | 64×64 | 33 | 4 | .6 | 180 |
| PKU | Siemens Trio 3T | 2000 | 30 | 90 | 210×210 | 64×64 | 30 | 4 | .8 | 210 |
| SCU | GE EXCITE 3T | 2000 | 30 | 90 | 220×220 | 64×64 | 30 | 5 | 0 | 200 |
| SWU | Siemens Trio 3T | 2000 | 30 | 90 | 220×220 | 64×64 | 32 | 3 | 1 | 242 |

Abbreviations: TR, repetition time; TE, echo time; FA, flip angle; FOV, field of view; CMU, China Medical University; CSU, Central South University; PKU, Peking University; SCU, Sichuan University; SWU, Southwest University; GE, General Electric.

#### China Medical University (CMU) dataset

R-fMRI images were acquired with a 3.0-T GE HDxT scanner (General Electric, Milwaukee, USA) with an 8-channel head coil. The parameters were TR=2,000 ms, TE=40 ms, flip angle=90°, field of view=240×240 mm^2^, and matrix=64×64. Thirty-five axial slices were collected with a 3-mm thickness and no gap. The scan lasted for 6 minutes and 40 s, resulting in 200 volumes.

#### Central South University (CSU) dataset

R-fMRI images were acquired with a 3.0-T GE HDxT scanner (General Electric, Milwaukee, USA) with a standard head coil. The parameters were TR=2,000 ms, TE=30 ms, flip angle=90°, field of view=220×220 mm^2^, and matrix=64×64. Thirty-three axial slices were collected with 4-mm thickness and a gap of 0.6 mm. The scan lasted for 6 minutes, resulting in 180 volumes.

#### Peking University (PKU) dataset

R-fMRI images were acquired with a 3.0-T Siemens Magnetom Trio scanner (Siemens, Erlangen, Germany) with a standard head coil. The parameters were TR=2,000 ms, TE=30 ms, flip angle=90°, field of view=210×210 mm^2^, and matrix=64×64. Thirty axial slices were collected with 4-mm thickness and a gap of 0.8 mm. The scan lasted for 7 minutes, resulting in 210 volumes.

#### Sichuan University (SCU) dataset

R-fMRI images were acquired with a 3.0-T GE EXCITE scanner (General Electric, Milwaukee, USA) with a standard head coil. The parameters were TR=2,000 ms, TE=30 ms, flip angle=90°, field of view=220×220 mm^2^, and matrix=64×64. Thirty axial slices were collected with a 5-mm thickness and no gap. The scan lasted for 6 minutes and 40 s, resulting in 200 volumes.

#### Southwest University (SWU) dataset

R-fMRI images were acquired with a 3.0-T Siemens Trio scanner (Siemens, Erlangen, Germany) with a 16-channel head coil. The parameters were TR=2,000 ms, TE=30 ms, flip angle=90°, field of view=220×220 mm^2^, and matrix=64×64. Thirty-two axial slices were collected with 3-mm thickness and a gap of 1 mm. The scan lasted for 8 minutes and 4 s, resulting in 242 volumes.

### Data Preprocessing

R-fMRI image preprocessing was conducted with SPM12 (www.fil.ion.ucl.ac.uk/spm/) and an in-house toolbox, SeeCAT (www.nitrc.org/projects/seecat). Briefly, the first ten time points (the first five time points for the CSU dataset due to the short scan time) were discarded. Subsequent preprocessing steps included slice-timing correction and head-motion correction. Next, motion-corrected functional images were normalized to the standard space using the EPI template, resampled to 3-mm isotropic voxels, and further smoothed with a 6-mm full-width at half maximum Gaussian kernel. Linear detrending was performed, and several confounding covariates, including the Friston-24 head-motion parameters, white matter and cerebrospinal fluid signals, were regressed out from the time series for all voxels. Subsequently, temporal bandpass filtering (0.01-0.1 Hz) was applied. Given that the spectral magnitudes in R-fMRI signals are sensitive to head motion (Satterthwaite et al., 2013), we also applied filtering with a band of 0.01-0.08 Hz for validation purposes. Finally, a “scrubbing” procedure was performed on individual preprocessed data to remove outlier data due to head motion (Power et al., 2012). Specifically, for volumes with a framewise displacement exceeding a threshold of 0.5 mm, we replaced the volumes and their adjacent volumes (2 forward and 1 backward frames) with linear interpolated data.

### Functional Brain Measurements

In the present study, we used three functional brain measurements, ALFF (Zang et al., 2007), ReHo (Zang et al., 2004) and distance-dependent FCS (Buckner et al., 2009; Dai et al., 2015; Liang et al., 2013; Liao et al., 2013; Xia et al., 2018), which have been widely used in previous R-fMRI studies in MDD. These three measures examine functional coordination ranging from focal activity to long-range connections, respectively. Specifically, i) the ALFF reflects functionally coordinated amplitudes among various neurons within a voxel (Zang et al., 2007). For a given gray matter (GM) voxel, the R-fMRI time course was first extracted and then converted to the frequency domain using a fast Fourier transform. The ALFF of this voxel was computed as the averaged square root of the power spectrum across the 0.01-0.1 Hz frequency interval. ii) The ReHo estimates functional similarities in brain activities among neighboring voxels located within a short range (Zang et al., 2004). For a given GM voxel, the ReHo value was computed as Kendall’s coefficient of concordance (Kendall and Gibbons, 1990) of the time series of this voxel with those of its nearest neighbors. iii) The FCS captures a total functional coordination between a given GM voxel and all other voxels at a specific distance range (Buckner et al., 2009; Dai et al., 2015; Wang et al., 2014a; Wang et al., 2015b; Xia et al., 2018). Briefly, for a given GM voxel, we first computed its full-range FCS by summing Pearson’s correlation coefficients between the voxel and other voxels. Considering the effects of distance on brain networks in healthy (Achard et al., 2006) and diseased (Alexander-Bloch et al., 2013; Dai et al., 2015; Wang et al., 2014b; Xia et al., 2018) populations, we further divided whole-brain functional connectivity into 9 bins with Euclidean distances binned into 20-mm steps, ranging from 0 to 180 mm (the longest distance between voxels in the GM mask), and calculated a distance-weighted FCS for each bin (Xia et al., 2018). Voxels with higher FCS values tend to play central roles in transferring information flow across regions. In the present study, we obtained individual ALFF, ReHo and FCS maps in a voxel-wise manner, which were further normalized to reduce global brain effects. Notably, all of the analyses were constrained within a GM mask that was generated by thresholding the GM probability map in SPM 12 with a threshold of 0.2 and removing voxels that were not covered by the subjects’ data. As a result, 12 individual functional brain maps (1 ALFF, 1 ReHo, 1 full-range FCS, and 9 distance-specific FCS maps) were generated for each subject, representing functional coordination from local voxels to long-range connectivity.

### Reproducibility Analyses of Functional Brain Measures

We systematically evaluated the effects of several methodological and clinical factors on the reproducibility of dysfunctions in MDD. These factors included the multisite statistical analysis strategies, removal of the global signal, across-center consistency, clinical variables (e.g., first episode, medication status, and age of onset) and sample size.

#### Multi-site statistical analysis strategies

To study whether MDD-related functional alterations can be reliably identified, we used two multisite statistical analysis methods, the stepwise linear regression and the Liptak-Stouffer *z*-score method (Liptak, 1958). i) Stepwise linear regression method. Briefly, we first pooled all individuals from the five centers together, and regressed out the center effect, and then established a stepwise linear regression model for each metric (i.e., ALFF, ReHo or FCS). In this model, the metric was treated as the dependent variable, and the age, sex, group, age-by-group, sex-by-group and age-by-sex-by-group interactions were treated as independent variables. For each voxel, the model started with the same variables, and the independent variables were iteratively removed from the model if they did not significantly predict the dependent variable, leading to a potentially unique final model. ii) Liptak-Stouffer *z*-score method (Liptak, 1958). This method has been used to analyze multisite MRI data (Cheng et al., 2016; Glahn et al., 2008). Briefly, for each metric, we first evaluated the between-group differences at each center by using a general linear model with the metric as a dependent variable, group as an independent variable, and age and sex as covariates. The *P*-value for the group effect derived from the GLM was first converted to its corresponding *z*-score, and a combined *Z*-score was then calculated as follows:

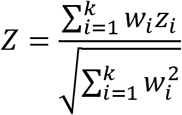

where *i*=1, 2, …, *k* represents the centers, 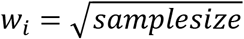 is the weight for the *i*th center, and *z_i_* is the *z*-score of the *i*th center. Finally, the *Z*-score was transformed to its corresponding *P*-value. Both statistical analysis strategies (i.e., stepwise linear regression and Liptak-Stouffer *z*-score method) were separately performed in a voxel-wise fashion, and the height threshold for significance was set at a two-tailed *P*<.001 at a voxel level to strictly control for the false positive rate (Eklund et al., 2016). Gaussian random-field correction at the cluster level was performed for multiple comparisons, with an extent significance level at *P*<.05.

#### Global signal processing strategies

The biological substrate of the global signal is currently unclear (Murphy and Fox, 2017), given that it simultaneously captures neural activity and physiological noise such as respiration and movement (Liu et al., 2017b; Murphy and Fox, 2017). To date, the procedure of performing global signal removal (GSR) or not during R-fMRI data preprocessing remains controversial. One recent study revealed the diverse effects of different processing strategies on global signals (with and without GSR) on functional data from subjects with psychiatric disorders (Yang et al., 2014). To estimate the effect of GSR on identifying MDD-related functional alterations, we reperformed the abovementioned two statistical strategies by analyzing R-fMRI data with GSR in preprocessing.

#### Cross-center consistency

We first assessed the site effect on these functional measurements and whether the multisite statistical analysis strategy of linear regression could reduce site effects. Briefly, for each functional metric, we performed Kruskal-Wallis tests across centers to estimate the site effect at each voxel. The obtained *P*-value map was converted to a *Z*-value map and corrected for multiple comparisons using Gaussian random-field correction. The significance level was set at a height threshold of two-tailed *P*<.001 with an extent threshold of *P*<.05. Then, we pooled all individuals from the five centers together and linearly regressed out the dummy site variables. The Kruskal-Wallis tests were performed again on the processed metric maps to check whether the site effects were reduced.

Additionally, we utilized a ComBat model to correct for the site effects (Yu et al., 2018). The ComBat model is based on multivariate linear mixed effects regression, in which the empirical Bayes is used to improve the estimation of the model parameters (Yu et al., 2018). We further used Kruskal-Wallis tests to estimate the site effects and performed the stepwise linear regression to identify MDD-related alterations in ComBat harmonized metric data.

Next, we conducted a conjunction analysis to assess the reproducibility of MDD-related functional alterations across different centers. Considering that the Liptak-Stouffer z-score method was performed based on statistical analysis in each center, which is conceptually similar to a conjunction method, we only performed this across-center consistency analysis for the stepwise regression method on both data with and without GSR. Briefly, for each identified cluster with a significant between-group difference, we first conducted the stepwise linear regression analysis in a voxelwise manner to obtain the group effect in each center, using the same model previously used in the pooled dataset. We then binarized the group effect maps in that voxels with a significant between-group difference were set as 1; otherwise, they were set as 0. Given the *post hoc* nature of this analysis, the significance level was set at *P*=.05 at the voxel level. Finally, the conjunction map for each cluster was obtained by summing the binarized group effect maps across centers.

#### Effect of clinical variables

We further investigated the effects of categorized clinical variables on the identification of group differences in functional activities. Briefly, we classified the patients into six different subgroups according to their clinical information, including patients in their first episode or in a nonfirst episodes, patients receiving medication or not receiving medication, and patients with an onset age older than or no more than 21 years (Benazzi, 2009; Schmaal et al., 2017; Schmaal et al., 2016). First, we performed statistical analysis on the clinical variables (i.e., illness duration, onset age, episode number, and HDRS) between each corresponding pair of the subgroups. Second, for each group, along with HC data, we extracted the mean value of functional metrics within each of the previously identified clusters and performed the stepwise linear regression analysis to estimate the group effect. Cohen’s *d* was also calculated for each cluster to show the effect size. Finally, we directly compared the mean functional metric of each region between each corresponding pair of subgroups using stepwise linear regression analysis. A threshold of false discovery rate (FDR) corrected *P*<.05 across regions was considered significant.

Next, for regions with significant group differences, we also explored the relationships between functional measurements and continuous clinical variables (i.e., HDRS, duration of illness, onset age and episode number). A robust regression analysis was performed with each clinical variable as the dependent variable, mean functional measurements in each cluster as the independent variable, and the center, age, and sex as covariance. We conducted this analysis in the pooled dataset and in each center to assess the reproducibility. The significance level was set to *P*<.05 with Bonferroni correction across different clusters.

#### Effect of sample size

To estimate the number of participants needed to identify MDD-related functional alterations, we performed a bootstrap simulation analysis. Briefly, we first randomly sampled subsets from the pooled cohort with different sample sizes (from 50 to 700 individuals in each group, with an interval of 50 individuals). Given that the between-group difference might derived from the center effect by unbiased sampling, we constrained our sampling procedure so that at least 50% of the individuals of the two groups should come from the same centers. Then, in each subset, we performed a stepwise regression analysis for each functional metric to identify regions with significant group effects. For each sample size, we conducted a random sampling and statistical analysis 1,000 times, and we defined the reproducibility rate at each voxel as the percentage of times that the given voxel exhibited a significant group difference in 1,000 simulations. Three significance levels (a height threshold of *P*<.001, *P*<.01, and *P*<.05 at a voxel level with an extent threshold of *P*<.05 estimated by using Gaussian random field at cluster level) were applied, respectively. Therefore, at each significance level, we obtained a curve of reproducibility rate against sample size at each voxel, and the smallest required sample size to reach a critical reproducibility rate of 50% was calculated. Finally, the smallest required sample size of each of the previously identified regions with significant MDD-related functional alterations was estimated as the minimum smallest sample size across all voxels within the cluster under reproducibility rate of 50%.

## Results

### Reproducible Functional Brain Alterations in Patients with MDD (Statistical Strategies and GSR Effects)

Figure 1 and Figure 2 show between-group difference maps of each functional measure (without thresholding). With a strict statistical analysis with multiple comparison corrections (a cluster-level corrected *P*<.05 with a voxel-level *P*<.001), we identified a repeated pattern of significant functional alteration in patients with MDD that was mainly distributed in the prefrontal, parietal and occipital regions regardless of multicenter statistical analysis strategies and whether the global signal was regressed (Figure 3). Specifically, patients with MDD had significantly lower functional activities in the right postcentral gyrus (PoCG, ReHo), the bilateral orbitofrontal cortices (OFC, FCS of 0-20 mm) and the bilateral middle and inferior occipital gyri (FCS of 60-80 and 80-100 mm) than did the HCs. Furthermore, patients with MDD exhibited significantly higher functional activities in the left triangular part of the inferior frontal gyrus (IFGtriang, ALFF), the right supramarginal gyrus (SMG, FCS of 0-20 and 20-40 mm), the bilateral precuneus (FCS of 80-100 mm), and the right superior frontal gyrus (SFG, FCS of 100-120 mm) than did the HCs (Table 3–6). We did not observe significant effects for age-by-group, sex-by-group, or age-by-sex-by-group interactions, although significant age and sex effects were observed for these functional measurements (Figure S1 and S2).

**Figure 1.**
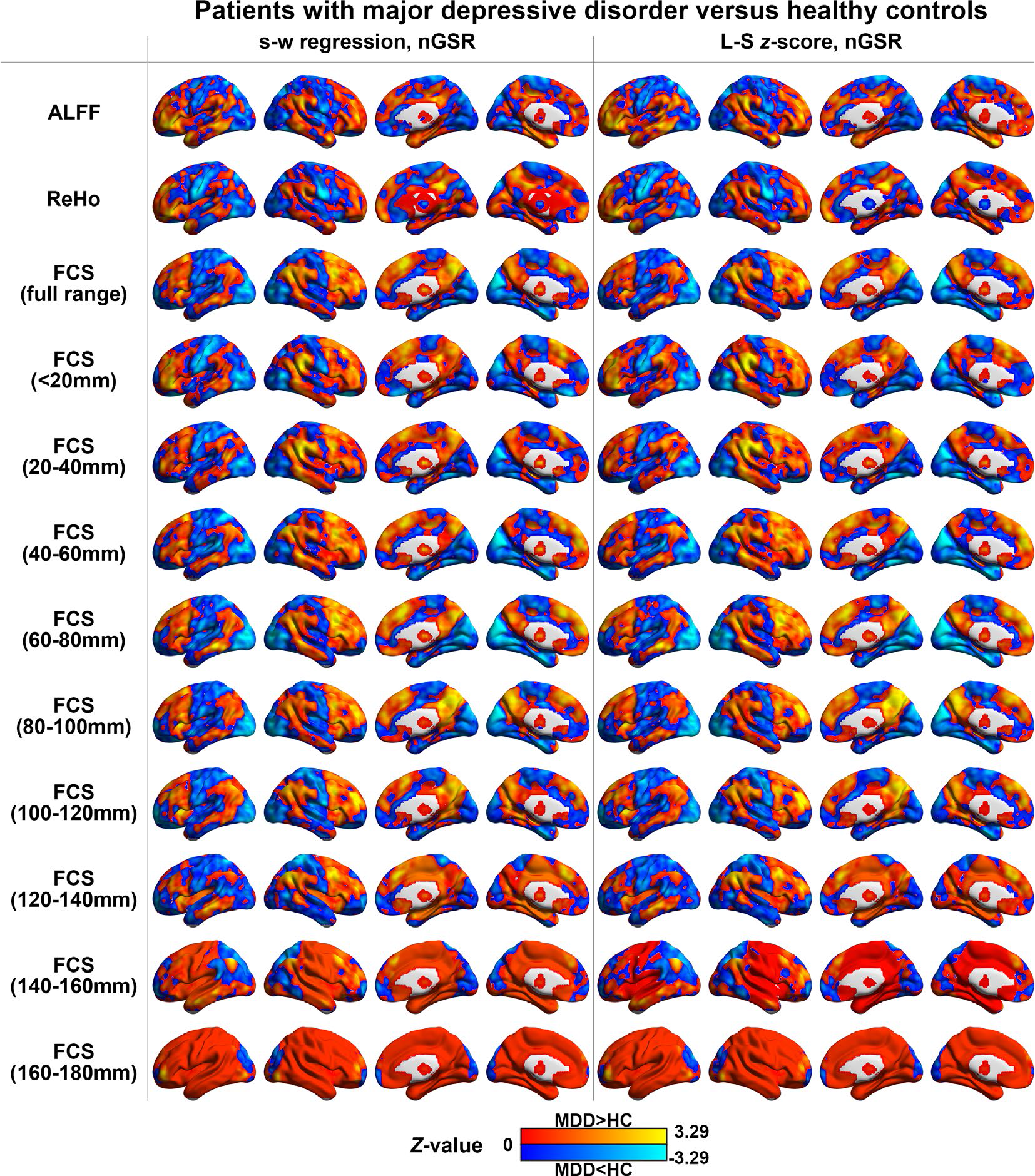
Differences in functional measurements between patients with MDD and healthy controls in data without GSR (not thresholding). The figure illustrates group effects on functional measures without thresholding using the stepwise regression analysis and Liptak-Stouffer z-score method in data without GSR. Warm and cold colors indicate higher and lower functional measurements in patients with MDD than in the HCs, respectively. MDD, major depressive disorder; HC, healthy controls; s-w, stepwise; nGSR, nonglobal signal regression; L-S, Liptak-Stouffer; ALFF, amplitude of low-frequency fluctuations; ReHo, regional homogeneity; FCS, functional connectivity strength.

**Figure 2.**
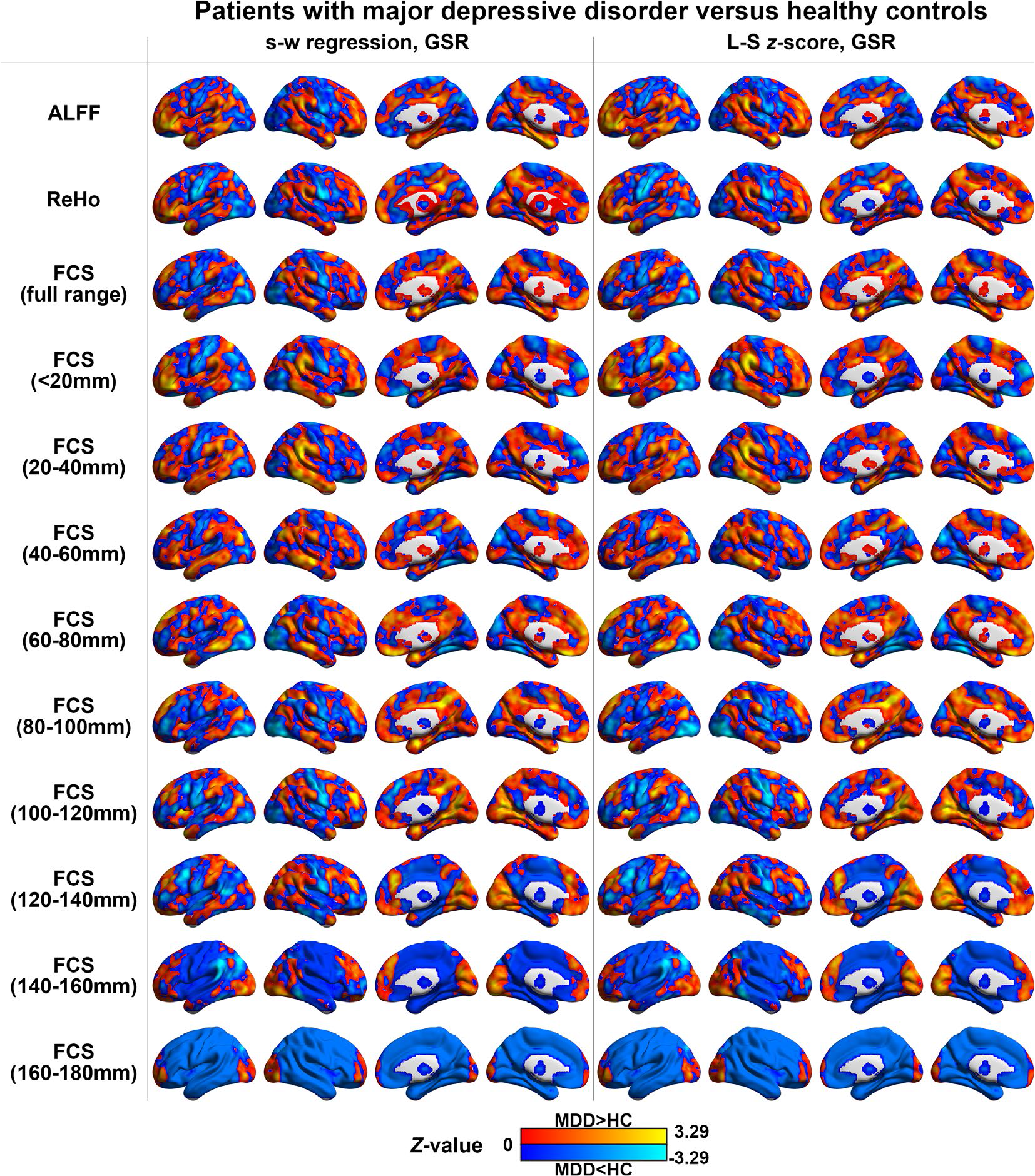
Differences in functional measurements between patients with MDD and healthy controls in data with GSR (not thresholding). The figure illustrates group effects on functional measures without thresholding using the stepwise regression analysis and Liptak-Stouffer z-score method in data with GSR. Warm and cold colors indicate higher and lower functional measurements in patients with MDD than in the HCs, respectively. MDD, major depressive disorder; HC, healthy controls; s-w, stepwise; L-S, Liptak-Stouffer; ALFF, amplitude of low-frequency fluctuations; ReHo, regional homogeneity; FCS, functional connectivity strength.

**Figure 3.**
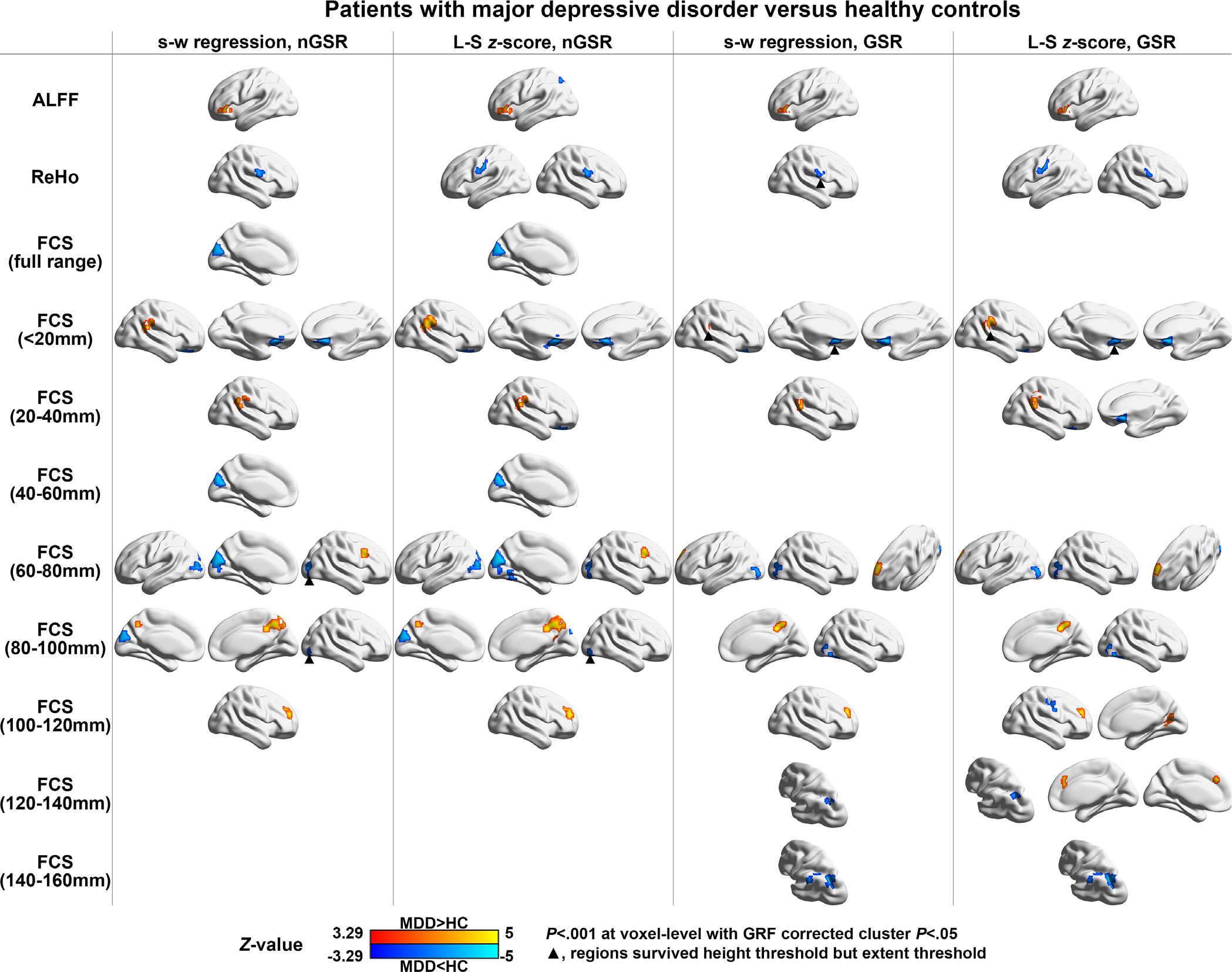
Differences in functional measurements between patients with MDD and healthy controls. The figure illustrates significant between-group effects on functional measurements from local voxel to distant brain connections using the stepwise regression analysis and Liptak-Stouffer *z*-score method on data with and without GSR. Warm and cold colors indicate higher and lower functional measurements in patients with MDD than in the HCs, respectively. The surface rendering was created using BrainNet Viewer (www.nitrc.org/projects/bnv/) (Xia et al., 2013). MDD, major depressive disorder; HC, healthy controls; s-w, stepwise; nGSR, nonglobal signal regression; L-S, Liptak-Stouffer; GSR, global signal regression; ALFF, amplitude of low-frequency fluctuations; ReHo, regional homogeneity; FCS, functional connectivity strength; GRF, Gaussian random field.

**Table 3.**
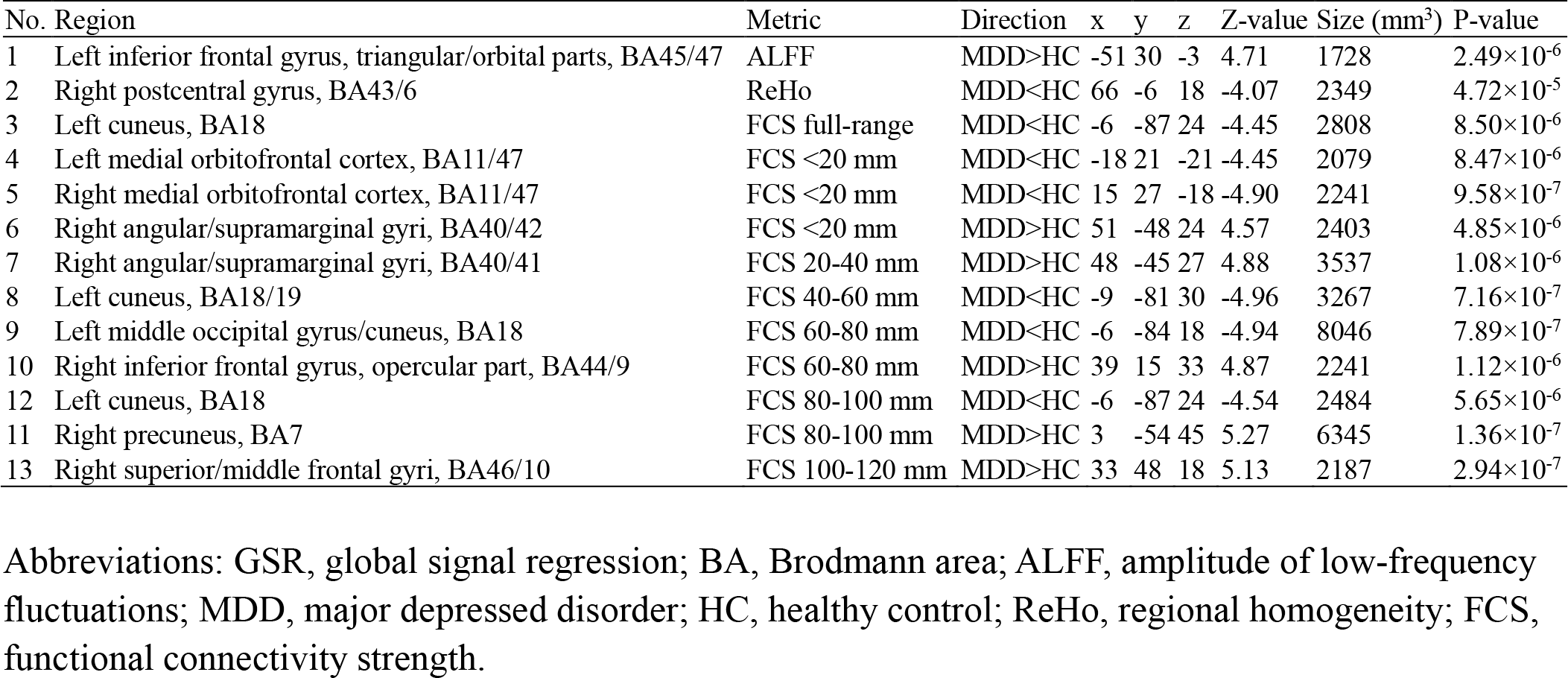
Clusters with significant group effects identified by stepwise regression analysis in data without GSR

| No. | Region | Metric | Direction | x | y | z | Z-value | Size (mm<sup>3</sup>) | P-value |
| --- | --- | --- | --- | --- | --- | --- | --- | --- | --- |
| 1 | Left inferior frontal gyrus, triangular/orbital parts, BA45/47 | ALFF | MDD>HC | -51 | 30 | -3 | 4.71 | 1728 | 2.49×10<sup>-6</sup> |
| 2 | Right postcentral gyrus, BA43/6 | ReHo | MDD<HC | 66 | -6 | 18 | -4.07 | 2349 | 4.72×10<sup>-5</sup> |
| 3 | Left cuneus, BA18 | FCS full-range | MDD<HC | -6 | -87 | 24 | -4.45 | 2808 | 8.50×10<sup>-6</sup> |
| 4 | Left medial orbitofrontal cortex, BA11/47 | FCS <20 mm | MDD<HC | -18 | 21 | -21 | -4.45 | 2079 | 8.47×10<sup>-6</sup> |
| 5 | Right medial orbitofrontal cortex, BA11/47 | FCS <20 mm | MDD<HC | 15 | 27 | -18 | -4.90 | 2241 | 9.58×10<sup>-7</sup> |
| 6 | Right angular/supramarginal gyri, BA40/42 | FCS <20 mm | MDD>HC | 51 | -48 | 24 | 4.57 | 2403 | 4.85×10<sup>-6</sup> |
| 7 | Right angular/supramarginal gyri, BA40/41 | FCS 20-40 mm | MDD>HC | 48 | -45 | 27 | 4.88 | 3537 | 1.08×10<sup>-6</sup> |
| 8 | Left cuneus, BA18/19 | FCS 40-60 mm | MDD<HC | -9 | -81 | 30 | -4.96 | 3267 | 7.16×10<sup>-7</sup> |
| 9 | Left middle occipital gyrus/cuneus, BA18 | FCS 60-80 mm | MDD<HC | -6 | -84 | 18 | -4.94 | 8046 | 7.89×10<sup>-7</sup> |
| 10 | Right inferior frontal gyrus, opercular part, BA44/9 | FCS 60-80 mm | MDD>HC | 39 | 15 | 33 | 4.87 | 2241 | 1.12×10<sup>-6</sup> |
| 12 | Left cuneus, BA18 | FCS 80-100 mm | MDD<HC | -6 | -87 | 24 | -4.54 | 2484 | 5.65×10<sup>-6</sup> |
| 11 | Right precuneus, BA7 | FCS 80-100 mm | MDD>HC | 3 | -54 | 45 | 5.27 | 6345 | 1.36×10<sup>-7</sup> |
| 13 | Right superior/middle frontal gyri, BA46/10 | FCS 100-120 mm | MDD>HC | 33 | 48 | 18 | 5.13 | 2187 | 2.94×10<sup>-7</sup> |

Abbreviations: GSR, global signal regression; BA, Brodmann area; ALFF, amplitude of low-frequency fluctuations; MDD, major depressed disorder; HC, healthy control; ReHo, regional homogeneity; FCS, functional connectivity strength.

The results from two different statistical analysis and GSR strategies were generally consistent. However, influences were observed for several specific regions. i) The Liptak-Stouffer *z*-score method was slightly more sensitive in identifying MDD-related alterations of a few regions with better extent significance, such as significantly lower ReHo in the left PoCG in MDD patients than in HCs (Figure 3; Table 3 vs. Table 4; Table 5 vs. Table 6). ii) In data without GSR, patients with MDD had significantly lower FCS in the left cuneus (CUN, full-range, 40-60, and 60-80 mm) and higher FCS in the right opercular part of the inferior frontal gyrus (IFGoperc, 60-80 mm) than did the HCs (Figure 3; Table 3 vs. Table 5; Table 4 vs. Table 6). In contrast, in data with GSR, we observed a significantly lower FCS in the right precentral gyrus (PrCG, 100-120 mm) and the left inferior parietal lobule (IPL, 120-140 and 140-160 mm), and a significantly higher FCS in the left SFG (60-80 mm), right calcarine fissure cortex (CAL, 100-120 mm), and the bilateral medial part of SFG (SFGmed,120-140 mm) in the MDD groups than in the HC group (Figure 3; Table 3 vs. Table 5; Table 4 vs. Table 6). These results suggest that the data after GSR were the most sensitive for identifying alterations in long-range functional coordination in patients with MDD. Finally, the results of using filtering with a band of 0.01-0.08 Hz during data preprocessing were overall parallel to the main findings in that most of the significant MDD-related alterations remains unchanged (Figure S3).

**Table 4.** Clusters with significant group effects identified by the Liptak-Stouffer z-score method in data without GSR

| No. | Region | Metric | Direction | x | y | z | Z-value | Size (mm<sup>3</sup>) | P-Value |
| --- | --- | --- | --- | --- | --- | --- | --- | --- | --- |
| 1 | Left inferior frontal gyrus, triangular/orbital parts, BA47 | ALFF | MDD>HC | -39 | 24 | -6 | 4.39 | 1917 | 1.144×10<sup>-5</sup> |
| 2 | Left superior parietal gyrus, BA7 | ALFF | MDD<HC | -12 | -81 | 45 | -4.42 | 1647 | 9.962×10<sup>-6</sup> |
| 3 | Right postcentral gyrus, BA43/6 | ReHo | MDD<HC | 66 | -6 | 18 | -4.28 | 2241 | 1.877×10<sup>-5</sup> |
| 4 | Left postcentral gyrus, BA43/6 | ReHo | MDD<HC | -60 | -6 | 30 | -4.58 | 2133 | 4.762×10<sup>-6</sup> |
| 5 | Left cuneus, BA18/19 | FCS full-range | MDD<HC | -6 | -87 | 24 | -4.57 | 3429 | 4.899×10<sup>-6</sup> |
| 6 | Left medial orbitofrontal cortex, BA11/47 | FCS <20 mm | MDD<HC | -21 | 18 | -21 | -4.77 | 2889 | 1.851×10<sup>-6</sup> |
| 7 | Right medial orbitofrontal cortex, BA11/47 | FCS <20 mm | MDD<HC | 18 | 21 | -21 | -4.95 | 2673 | 7.421×10<sup>-7</sup> |
| 8 | Right angular/supramarginal gyri, BA40/42 | FCS <20 mm | MDD>HC | 51 | -45 | 24 | 4.89 | 5940 | 1.008×10<sup>-6</sup> |
| 9 | Right angular/supramarginal gyri, BA40/41 | FCS 20-40 mm | MDD>HC | 48 | -45 | 27 | 5.07 | 5508 | 4.084×10<sup>-7</sup> |
| 10 | Left cuneus, BA18/19 | FCS 40-60 mm | MDD<HC | -6 | -75 | 21 | -5.57 | 7344 | 2.518×10<sup>-8</sup> |
| 11 | Right middle occipital gyrus, BA19/18 | FCS 60-80 mm | MDD<HC | 36 | -87 | 15 | -4.07 | 2457 | 4.641×10<sup>-5</sup> |
| 12 | Left middle occipital gyrus/cuneus, BA18/19 | FCS 60-80 mm | MDD<HC | -6 | -84 | 18 | -4.94 | 8046 | 7.893×10<sup>-7</sup> |
| 11 | Right inferior frontal gyrus, opercular part, BA44/9 | FCS 60-80 mm | MDD>HC | 39 | 15 | 33 | 5.16 | 2403 | 2.430×10<sup>-7</sup> |
| 14 | Left cuneus, BA18/19 | FCS 80-100 mm | MDD<HC | -6 | -84 | 21 | -4.77 | 3402 | 1.806×10<sup>-6</sup> |
| 15 | Right precuneus, BA7 | FCS 80-100 mm | MDD>HC | 6 | -54 | 45 | 5.44 | 9288 | 5.269×10<sup>-8</sup> |
| 16 | Right superior/middle frontal gyri, BA46/10 | FCS 100-120 mm | MDD>HC | 33 | 48 | 18 | 5.50 | 3969 | 3.820×10<sup>-8</sup> |

Abbreviations: GSR, global signal regression; ALFF, amplitude of low-frequency fluctuations; MDD, major depressed disorder; HC, healthy control; ReHo, regional homogeneity; FCS, functional connectivity strength.

**Table 5.** Clusters with significant group effects identified by stepwise regression analysis in data with GSR

| No. | Region | Metric | Direction | x | y | z | Z-value | Size (mm<sup>3</sup>) | P-Value |
| --- | --- | --- | --- | --- | --- | --- | --- | --- | --- |
| 1 | Left inferior frontal gyrus, triangular/orbital parts, BA45/47 | ALFF | MDD>HC | -51 | 30 | -3 | 4.66 | 1377 | 3.240×10<sup>-6</sup> |
| 2 | Right medial orbitofrontal cortex, BA11/47 | FCS <20 mm | MDD<HC | 15 | 27 | -18 | -4.87 | 2916 | 1.139×10<sup>-6</sup> |
| 3 | Right supramarginal gyrus, BA40 | FCS 20-40 mm | MDD>HC | 51 | -42 | 24 | 4.76 | 2619 | 1.898×10<sup>-6</sup> |
| 4 | Left middle occipital gyrus, BA19/18 | FCS 60-80 mm | MDD<HC | -39 | -87 | 3 | -4.31 | 1269 | 1.603×10<sup>-5</sup> |
| 5 | Right middle occipital gyrus, BA19/18 | FCS 60-80 mm | MDD<HC | 42 | -81 | 6 | -4.11 | 1566 | 3.889×10<sup>-5</sup> |
| 6 | Left superior frontal gyrus, BA9 | FCS 60-80 mm | MDD>HC | -15 | 57 | 33 | 5.66 | 1674 | 1.540×10<sup>-8</sup> |
| 7 | Right inferior occipital gyrus, BA19 | FCS 80-100 mm | MDD<HC | 39 | -72 | -12 | -4.30 | 1539 | 1.685×10<sup>-5</sup> |
| 8 | Right precuneus, BA7 | FCS 80-100 mm | MDD>HC | 9 | -45 | 39 | 5.23 | 2349 | 1.668×10<sup>-7</sup> |
| 9 | Right superior/middle frontal gyri, BA46/10 | FCS 100-120 mm | MDD>HC | 27 | 45 | 21 | 4.93 | 1890 | 8.393×10<sup>-7</sup> |
| 10 | Left angular gyrus/inferior parietal lobule, BA40/7 | FCS 120-140 mm | MDD<HC | -33 | -51 | 36 | -4.49 | 1350 | 7.122×10<sup>-6</sup> |
| 11 | Left inferior parietal lobule, BA40 | FCS 140-160 mm | MDD<HC | -48 | -45 | 36 | -5.44 | 5967 | 5.480×10<sup>-8</sup> |

Abbreviations: ALFF, amplitude of low-frequency fluctuations; MDD, major depressed disorder; HC, healthy control; ReHo, regional homogeneity; FCS, functional connectivity strength.

**Table 6.**
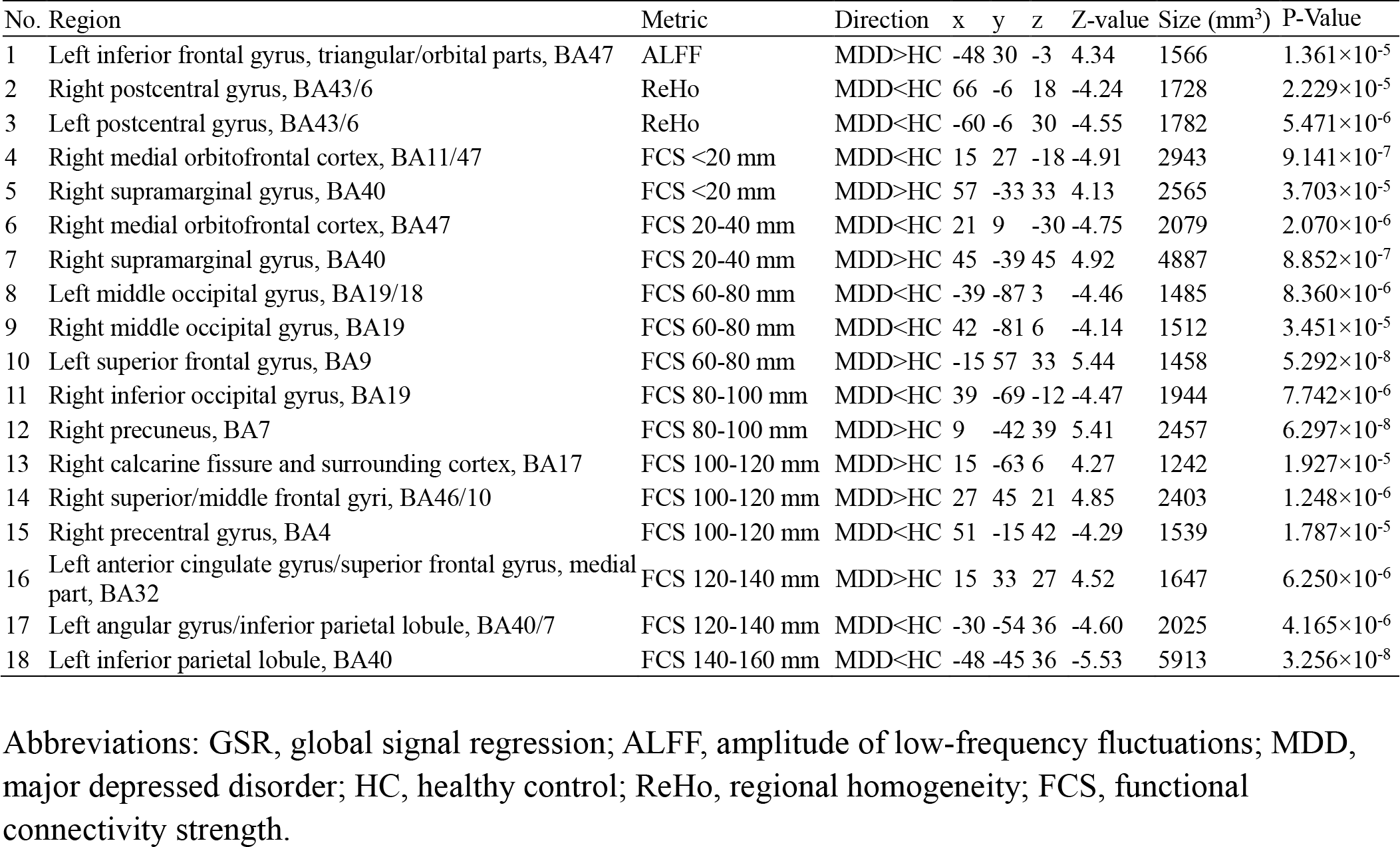
Clusters with significant group effects identified by the Liptak-Stouffer z-score method in data with GSR

### The Consistency of Cross-center Validation

We observed a significant site effect on all of the raw functional metrics in most of the brain regions (Figure S4), suggesting the necessity for performing site effect correction in multicenter imaging studies. Notably, these site effects no longer existed after performing linear regression or applying the ComBat method (except for ALFF in a few parietal and temporal regions in linear regressed maps, which did not overlap with regions exhibiting MDD-related alterations) (Figure S4). The result indicated the ability of both methods to correct confounding site factors in the MDD R-fMRI studies. Additionally, almost all of the MDD-related functional alterations remained significant in data processed with ComBat method (Figure S5).

Figure 4 illustrates the conjunction maps, showing significant between-group differences using stepwise regression analysis across five centers. Overall, we found a fair-to-good across-center consistency for these MDD-related functional alterations in both data with and data without GSR. Specifically, in data without GSR, the between-group differences in the left IFGtriang (ALFF), right SMG (FCS of 20-40 mm), left CUN (FCS of 40-60 mm), and PCUN (FCS of 100-120 mm) could be reproduced in four centers, while those in the other regions were observed three times. In data with GSR, alterations in regions located in the frontal and parietal cortices were more repeatedly identified (four times) across centers, including the right SMG (FCS of 20-40 mm), left SFG (FCS of 60-80 mm), PCUN (FCS of 100-120 mm), right SFG (FCS of 100-120 mm), and left IPL (FCS of 140-160 mm).

**Figure 4.**
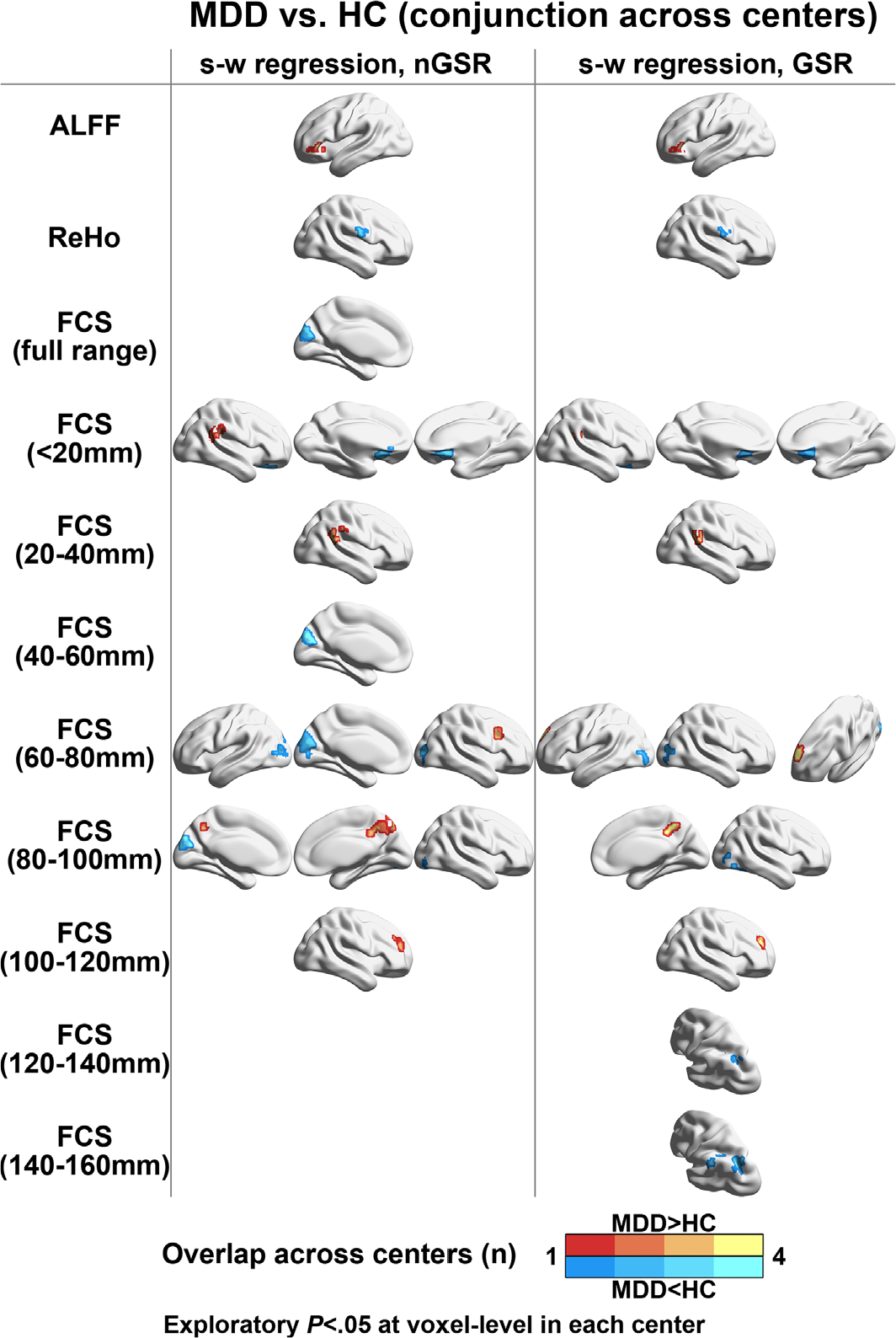
Conjunction maps across centers on functional differences between MDD and control groups. The figure illustrates the number of centers showing significant between-group differences within each cluster across five centers. Warm and cold colors indicate higher and lower functional measurements in patients with MDD than in the HCs, respectively. MDD, major depressive disorder; HC, healthy controls; s-w, stepwise; nGSR, nonglobal signal regression; GSR, global signal regression; ALFF, amplitude of low-frequency fluctuations; ReHo, regional homogeneity; FCS, functional connectivity strength.

### Effects of First Episode, Medication Status, and Onset Age

Regarding clinical variables, the first-episode patients had a significantly shorter illness duration and a lower rate of receiving medication than did the recurrent patients (both *P*<.001). Patients with medication had significantly longer illness duration, higher episode number, lower HDRS, and lower first-episode ratio than patients that did not receive medication (all *P*<.001). Moreover, there were no significant differences in illness duration, episode number, HDRS, or medication status between patients with an onset age greater than or no more than 21 years (Table S1).

For the functional brain alterations, after dividing the patients into first-episode and recurrent groups, all identified regions remained significant in the first-episode patients and mostly in the recurrent patients (all *P*<.05, FDR corrected), except for several regions with short-range functional coordination, such as the left IFGtriang, right PoCG, bilateral OFC, and left cuneus (Figure 5). After dividing the patients into medicated and nonmedicated groups, the between-group differences for all the clusters remained significant in both groups (all *P*<.05, FDR corrected) (Figure 5). After dividing patients according to the onset age, patients with MDD who had an onset age greater than 21 years showed short-range functional alterations (ALFF, ReHo, and FCS of <60 mm), including the left IFGtriang, right PoCG, right SMG, left OFC and left cuneus. In contrast, patients with MDD who had an onset age no greater than 21 years mainly exhibited altered long-range functional coordination (FCS of >60 mm), including the right IFGoperc, bilateral cuneus, bilateral precuneus and right SFG (all *P*<.05, FDR corrected) (Figure 5). Finally, in the direct comparisons between each corresponding pair of patient subgroups, we only found significantly lower FCS (80-100 mm) of the left cuneus in patients with an onset age no greater than 21 years than in patients with an onset age greater than 21 years (*P*=.004, FDR corrected). No significant differences were observed between first-episode and recurrent patients or between patients with and without medication.

**Figure 5.**
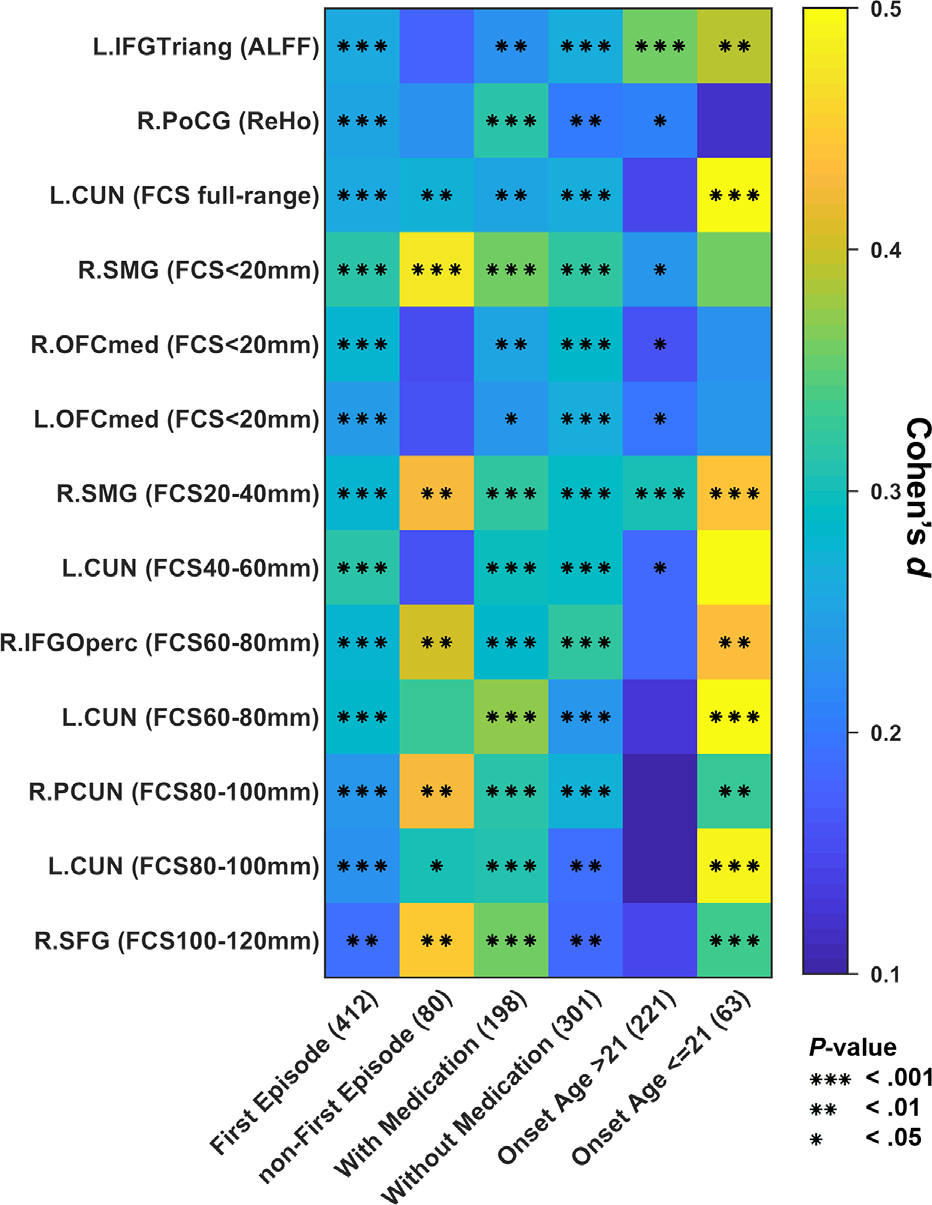
Effects of clinical variables on clusters showing significant between-group differences. The number of each clinical variable represents the number of patients in each group. Mean Cohen’s *d* and *P*-values were calculated across the voxels within each cluster showing significant between-group differences identified in all patients. ALFF, amplitude of low-frequency fluctuations; ReHo, regional homogeneity; FCS, functional connectivity strength; IFGTriang, inferior frontal gyrus, triangular part; PoCG, postcentral gyrus; SMG, supramarginal gyrus; OFCmed, medial orbitofrontal cortex; CUN, cuneus; IFGOperc, inferior frontal gyrus, opercular part; PCUN, precuneus; SFG, superior frontal gyrus.

### Correlations Between Functional Metrics and Clinical Variables in Patients with MDD

In the pooled dataset, we observed a significant positive correlation between the long-range FCS of the right PCUN (80-100 mm) and episode number in patients with MDD (*P*=.003, Bonferroni-corrected) (Figure 6). Additionally, we observed a negative correlation between the ReHo of the right PoCG and the HDRS (*P*=.021, uncorrected) (Figure 6). None of the functional metrics were correlated with the duration of illness or age of onset. In the dataset of each center, we found a positive correlation between the FCS of the right IFGOperc (60-80 mm) and the illness duration in the CSU dataset (*P*=.009, uncorrected); a negative correlation between the FCS of the right SMG (20-40 mm) and the age of onset in the CSU dataset (*P*=.038, uncorrected); a negative correlation between the ReHo of the right PoCG and the age of onset in the SCU dataset (*P*=.034, uncorrected); and a negative correlation between the ReHo of the right PoCG and the HDRS in the CMU dataset (*P*=.010, uncorrected) (Figure S6 and Table 7). These results indicate poor cross-center reproducibility of the relationship between functional metrics and clinical variables.

**Table 7.** Relationship between functional measurements and clinical variables

| Dataset | Region | Metric | Clinical variables | T-value | P-Value | Bonferroni Corrected P-value |
| --- | --- | --- | --- | --- | --- | --- |
| Pooled | Right precuneus | FCS 80-100 mm | Episode number | 3.02 | .003 | .037 |
| Pooled | Right postcentral gyrus | ReHo | HDRS-17 | -2.32 | .021 | .270 |
| CSU | Right inferior frontal gyrus, opercular part | FCS 60-80 mm | Illness duration | 2.66 | .009 | .111 |
| CSU | Right supramarginal gyrus | FCS 20-40 mm | Onset age | -2.10 | .038 | .499 |
| SCU | Right postcentral gyrus | ReHo | Onset age | -2.24 | .034 | .442 |
| CMU | Right postcentral gyrus | ReHo | HDRS-17 | -2.63 | .010 | .125 |

Abbreviations: FCS, functional connectivity strength; ReHo, regional homogeneity; HDRS, Hamilton depression rating scale; CMU, China Medical University; CSU, Central South University; SCU, Sichuan University.

**Figure 6.**
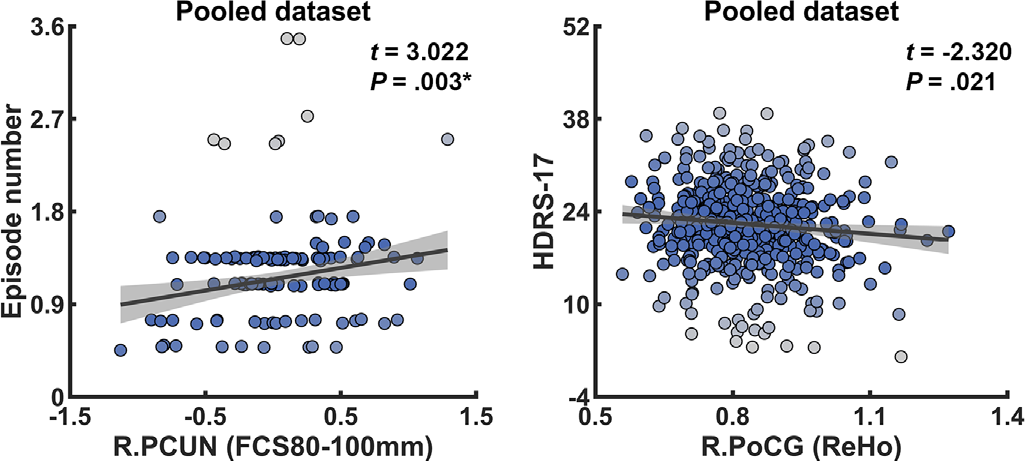
Robust fitting of functional measurements and clinical variables in pooled dataset. Each dot represents a subject, and its color indicates its weight in the robust regression analysis. A color map of blue to gray indicates the regression weight from high to low, respectively. Dashed lines indicate the confidence interval of the regression. The functional measurements and clinical variables were fitted by age, sex and centers. PCUN, precuneus; PoCG, postcentral gyrus; HDRS, Hamilton depression rating scale; FCS, functional connectivity strength.

### Influence of Sample Size on Reproducibility

At a cluster level (for previously identified cluster in Table 3), a sample size of 400 to 700 was needed to reach a reproducibility rate of 50% for a corrected *P*<.05 with a height threshold of *P*<.001 (Figure 7A, left). The required sample size depends on different regions and functional measurements. Notably, the between-group differences in long-range FCS required relatively a smaller sample size than did those in short-range FCS, ReHo, and ALFF of the regions identified to be reproduced in the same chance. Of the functional measurements, the FCS of the left CUN (40-60 and 60-80 mm) and the right PCUN (80-100 mm) were the most reproducible regions, requiring 400 to 475 subjects in each group to show corrected between-group differences with a chance of 50%. The ReHo in the right PoCG was the least reproducible region, and even with 700 subjects in each group, the chance for surviving correction was less than 50%. With the reduction of height significance level, the overall pattern of curves among different clusters remain unchanged but with smaller sample size required. The FCS of the left CUN (60-80 mm) and the right PCUN (80-100 mm) were still the most reproducible regions, requiring 300 and 225 for a height threshold of *P*<.01 and *P*<.05, respectively (Figure 7A, middle and right). Additionally, the reproducibility rate of several regions dropped under these lenient thresholds even when sample increased, including the ReHo of the right PoCG, the FCS of the right SMG and left OFC (<20 mm), and the FCS of the right SFG (100-120 mm). These might be largely due to the limited size of these significant clusters that cannot survive the corrections of large extent thresholds estimated with low height thresholds. Figure 7B presents the required sample size at a voxel level under different significance levels.

**Figure 7.**
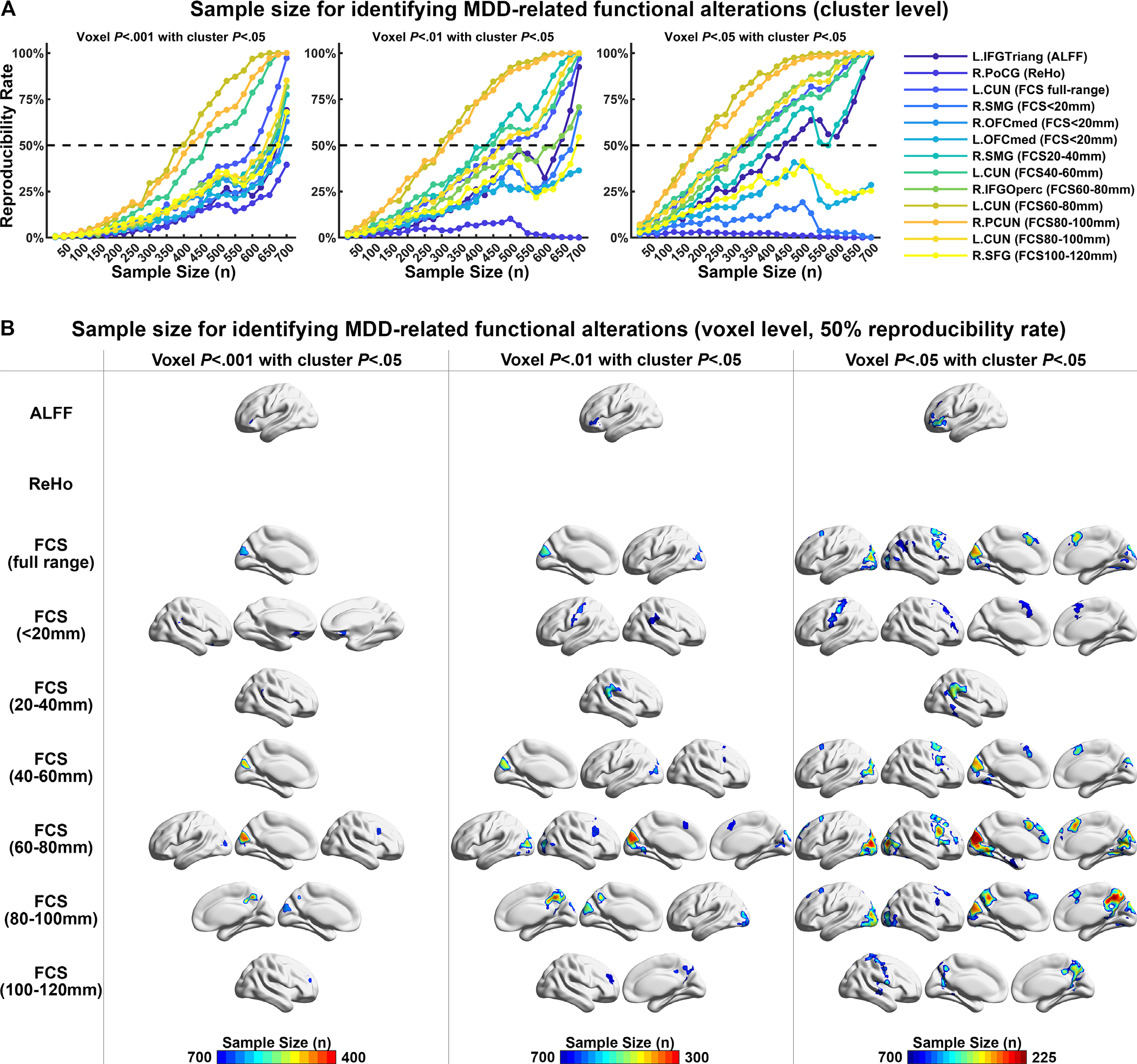
Influence of sample size on reproducibility rate. (A) This figure illustrates the relationship between the sample size in each group and the reproducibility rate of identifying each region with significant between-group differences. Each dot indicates the percentile in 1,000 bootstrapping simulations of n subjects in each group. (B) This figure shows brain maps of the least number of participants in each group required to reach a reproducibility rate of 50% for identifying significant between-group differences at different significance levels. The reproducibility rate for each voxel was estimated as the percentage of times with significance in 1,000 bootstrapping simulations.

## Discussion

Using a large-sample, multicenter R-fMRI dataset from a Chinese MDD cohort (N = 1,434), we revealed a repeated pattern of hypoactivity in the orbitofrontal, sensorimotor and visual cortices and hyperactivities in the frontoparietal cortices in MDD patients compared to HCs. These findings were generally reproducible regardless of statistical strategies, removal of global signal and medication status, and exhibited a high across-center consistency. However, the between-group differences were partially influenced by the episode status and the age of onset in patients, and the correlations between functional metric and clinical variables had poor across-center reproducibility. Finally, we showed that at least 400 subjects in each group were required to replicate significant alterations (a voxel level of *P*<.001 and a cluster level of *P*<.05) at a 50% reproducible chance. Together, these results highlight reproducible patterns of functional alterations in MDD and influencing factors involving research methodology and clinical design.

### MDD-relevant Hypoactivity in the Medial Orbitofrontal and Primary Cortices

The OFC is involved in prediction and decision making regarding emotion-related information for hedonic experiences (Kringelbach, 2005). Particularly, the medial OFC is associated with reward processing, including reward reinforcement, learning and memory, and is a crucial hub in the reward circuit connecting the medial temporal lobe and prefrontal cortex (Kringelbach and Rolls, 2004; Rolls, 2016). Previous studies in MDD reported a wide range of abnormalities in this region in either structure or function, such as reduced GM volume (Bremner et al., 2002) and cortical thickness (Schmaal et al., 2017) and abnormal functional activation (Lawrence et al., 2004) and connectivity (Cheng et al., 2016; Meng et al., 2014; Wang et al., 2014a; Zhang et al., 2011). Here, our findings of functional disruption in the medial OFC provide further evidence of altered memory systems encoding pleasant feelings and rewards that underlie the persistently depressed mood or loss of interest in activities in MDD patients. Significant hypoactivity of the right opercular part of the PoCG and the cuneus was also observed in patients with MDD. Neuroimaging studies suggest that the opercular part of the PoCG is a functionally heterogeneous region involved in recognizing emotions from visually presented facial expressions (Adolphs et al., 2000). Although several studies observed MDD-related structural or functional abnormalities in the PoCG (Iwabuchi et al., 2015; Schmaal et al., 2017), disruptions in this specific location have been reported only rarely in patients with MDD. In contrast, abnormalities in the visual cortex, such as a reduced surface area (Schmaal et al., 2017) and decreased cerebral blood flow (Ito et al., 1996), are associated with depression. Moreover, connectome studies have also revealed disrupted network topologies in either functional or structural brain networks of the visual cortex in patients with MDD (Korgaonkar et al., 2014; Singh et al., 2013; Zhang et al., 2011). Interestingly, the orbitofrontal and primary cortices had a large effect size on the cortical area shrinkage observed in the large-sample, worldwide brain structural study performed by the ENIGMA consortium (Schmaal et al., 2017). This finding indicates common disruptions in structure and function of these brain regions in MDD.

### MDD-relevant Hyperactivity in the Frontoparietal Cortices

We observed that patients with MDD exhibited significant hyperactivity in the left IFGtriang, right SMG, right IFGoperc, bilateral precuneus and right SFG. MDD-related changes in these regions have been reported in several previous studies. For instance, higher glucose metabolism rates in the IFGtriang were observed in depressed patients (Biver et al., 1994), and normalized metabolism of this region was lower in paroxetine responders than in nonresponders (Brody et al., 1999). Using R-fMRI, Zhang *et al*. revealed greater nodal centralities of the right SMG and the right IFGoperc in drug-naïve patients with first-episode MDD, suggesting more involvement of these areas in coordinating whole-brain functional networks (Zhang et al., 2011). In patients with depression, the precuneus shows multidimensional hyperactivity, including increases in cerebral metabolism (Smith et al., 2009), within-system coordination (Greicius et al., 2007; Sheline et al., 2010), and functional network connectivity (Cheng et al., 2016; Zhang et al., 2011). The SFG is also an area that is deeply involved in the pathology of MDD, with higher regional glucose metabolism (Brody et al., 2001) and hyper intra- and intersystem coordination (Sheline et al., 2010). To a certain extent, these regions share a common feature: involvement in nonreward, emotion-related processing. Specifically, the IFGtriang is related to lexical-semantic processing (Gold et al., 2006) and the self-regulation of emotions (Johnston et al., 2010). The right supramarginal gyrus participates in cognitive or emotional processing series, including working memory (Liu et al., 2017a) and empathic judgments (Silani et al., 2013). The right IFGoperc, a homologous region of Broca’s area in the opposite hemisphere, could be associated with orthography-to-semantic (Siok et al., 2004) and perception of negative mood (Hofer et al., 2006). The precuneus is a highly heterogeneous region involved in a wide spectrum of integrated functions, including memory retrieval, self-consciousness, and emotion judgment (Cavanna and Trimble, 2006). The precuneus is also a core component of the default-mode network (Raichle et al., 2001) and a critical hub with dense, long-range connections in human whole-brain structural (Gong et al., 2009; van den Heuvel and Sporns, 2011) and functional (Buckner et al., 2009; Liang et al., 2013) networks. Based on these findings, the precuneus plays an important role in integrating brain functions. Furthermore, the SFG is another important cortical area responsible for a series of high-order cognitive functions, such as working memory (Liu et al., 2017a), moral decision making (Greene et al., 2001), and behavioral inhibition (Aron et al., 2004). Recent functional imaging studies have also revealed the recruitment of the SFG during the regulation of negative emotions through reappraisal/suppression strategies (Levesque et al., 2003; Phan et al., 2005). Together, the hyperactivity in these key brain areas contributes to the broad spectrum of emotion-related disturbances and cognitive deficits observed in subjects with depression.

### The Effect of Global Signal on Reproducibility of MDD-relevant Functional Alterations

The global signal of fMRI data refers to the averaged time course across all voxels of the brain areas, and its biological significance relevant to neuronal activity is still poorly understood (Murphy and Fox, 2017). Recent studies documented that the global signal is a complex mixed signal because it simultaneously captures the underlying neural activity (Scholvinck et al., 2010) and several confounding noises, such as motion, cardiac and respiratory cycles that are globally embedded in the fMRI signals in nature (Liu et al., 2017b; Murphy et al., 2013). Therefore, whether to perform global signal regression remains confusing and controversial because the conduction of GSR can not only, at least partially, reduce the effect of unnecessary global confounds and enhance the identification of system-specific connections (Fox et al., 2009), but also can partly wipe away potential personal traits or diagnostic information (Liu et al., 2017b) and mathematically mandates anti-correlations between regions (Murphy et al., 2009). GSR also affects the reliability of commonly used functional metrics, such as ReHo (Zuo et al., 2013) and functional connectivity (Chai et al., 2012), as well as the topology of functional networks (Chen et al., 2018; Liang et al., 2012; Tomasi et al., 2016). More importantly, the relationship between global signal and spatially distributed brain regions was altered in several neuropsychiatric disorders, such as schizophrenia (Yang et al., 2017) and autism spectrum disorder (Gotts et al., 2012), and the conduction of GSR affected the identification of functional alterations in schizophrenia (Yang et al., 2014) and Alzheimer’s disease (Chen et al., 2018). Here, we showed that the MDD-relevant alterations were largely reproducible in data with and without GSR for local functional metrics (i.e., ALFF and ReHo) and short-range FCS, while GSR enhanced the sensitivity in detecting alterations in long-range FCS. A possible reason is the shifting of resting-state functional correlation distribution after GSR from being mainly positive to being zero centered (Chai et al., 2012). In functional connections with a relatively high strength (e.g., >.2), the proportion of medium- and short-range connections is much larger than that of long-range connections. Therefore, the distribution of functional correlation mainly reflects the strength of medium and short-range connections, and the determination of whether conducting GSR could have little effect on changing the shape of the overall distribution. Conversely, given the small number of long-range functional connections, their strength distribution is much easier to influence by GSR, and thus might increase the sensitivity to identify disorder-relevant alterations.

### Advantages of a Large-sample Dataset

Our bootstrapping simulation study showed that at least a sample size of 400 subjects in each group was required to replicate significant MDD-relevant functional alterations with a reproducibility rate over 50%, and the minimum sample size largely depended on brain regions and significance thresholds. This simulation analysis might partially explain the large discrepancies in MDD-relevant functional alterations across previous R-fMRI studies. To our knowledge, this study represents the largest R-fMRI MDD dataset collected in China. One thousand four hundred thirty-four patients with MDD and HCs were included based on the consistent diagnostic standards of the DSM-IV for MDD. Although the patients were recruited from five different research centers (i.e., Changsha, Beijing, Chongqing, Shenyang, and Chengdu), this cohort exhibited remarkable homogeneity in biological and demographic characteristics, including genetic background, language, and cultural environment. This homogeneity enables the reliable and reproducible identification of key brain nodes with abnormal functional activities in Chinese patients with MDD due to improved cross-validated power in statistical analysis and the avoidance of biased sampling in small datasets. The promising international multicenter structural imaging studies conducted by the ENIGMA consortium (Schmaal et al., 2017; Schmaal et al., 2016) have provided an important analytical framework for delving into neuroimaging big data and have revealed the general, reliable structural underpinnings of MDD, regardless of racial, genetic and environmental factors. Together, these large-sample multisite studies are particularly helpful for identifying reliable brain nodes with MDD-relevant abnormalities and developing imaging biomarkers for an effective early diagnosis and optimization of interventions for patients with MDD (Drysdale et al., 2017; Perrin et al., 2012). It should also be noted that given the existence of large individual heterogeneity in the brain relative to a normative distribution, particularly in patients with psychiatric disorders (Marquand et al., 2016; Wolfers et al., 2018), these functional and structural alterations in MDD should be considered a general pattern of this disorder. The assessment of reproducibility of functional alterations at an individual level is highly encouraged and could improve understanding of individualized precision medication for depression.

### Limitations and Further Considerations

Several issues need to be addressed. First, the clinical information, such as medication status and episode number, was not fully recorded for each patient due to variations in data management practices across different centers. This issue limited our power to analyze the effects of clinical variables on functional alterations. Second, significant site effects were observed on the raw functional metrics, which might result from the different MRI scanners, imaging parameters, experimental procedures, and clinical status of patients across different centers. Although our statistical analysis strategy eliminated the linear effects of centers, the potential nonlinear and interaction effects of these factors might remain upon identifying MDD-related functional alterations. The heterogeneity in collecting clinical and imaging data could explain the poor reproducibility of correlations between functional metrics and clinical variables. Third, longitudinal datasets were not included in the current study. Future studies including longitudinal neuroimaging data will be critical for evaluating the reproducibility of probing mechanisms of the progression of depression in the brain and developing reliable biomarkers for early diagnosis and prediction of treatment effects. Fourth, a recent study offered a promising example of defining subtypes of depression using big imaging data (Drysdale et al., 2017). Although MDD subtypes are not the main topic of the current paper, classifying subtypes with multicenter datasets is an intriguing challenge when facing the high heterogeneity in data across centers. Constructing an appropriate cross-center normative modal and further assessing the reproducibility of the shared and distinct patterns of alterations in patients with different subtypes of MDD will benefit the diagnosis and optimization of treatment for affected individuals. Fifth, our study identified several repeatable key brain regions with significantly altered functional activities; however, we have not determined whether these alterations are coupled with underlying structural and metabolic substrates. Sixth, the FCS used in the current study is an integration functional metric that sums many of region-to-region connections. Investigations focusing on the reproducibility of MDD-related alterations in specific regional connections can provide insights into the understanding of crucial pathways in depression. Finally, depression is highly associated with various types of cognitive deficits and genetic risk factors, such as 5-HTTLPR (Caspi et al., 2003). Future studies combining neuroimaging data and these multidimensional biological variables will contribute to improving our understanding of the biological mechanisms of behavioral disturbance. Moreover, these studies provide crucial opportunities to probe the intermediate effects of genes and behavior on depression pathology in the brain and the development of high-dimensional biomarkers for individuals with depression.

