## Supplementary Information for "Reproducibility of functional brain alterations in major depressive disorder: evidence from a multisite resting-state functional MRI study with 1,434 individuals"

### Participants

A total of 1,558 participants, including 782 patients with major depressive disorder (MDD) and 776 healthy controls (HCs), were recruited from 5 research centers in China (The Second Xiangya Hospital, Central South University, CSU; Peking University Sixth Hospital, PKU; First Affiliated Hospital of Chongqing Medical School in Chongqing, CMS; First Affiliated Hospital of China Medical University, CMU; and West China Hospital of Sichuan University, SCU). The detailed inclusion and exclusion criteria for each center are listed below.

***CMU dataset:*** Four hundred thirty-one participants were recruited for participation in this study, including 155 patients with MDD and 276 HCs. All patients with MDD were recruited from the outpatient clinic at the Department of Psychiatry, First Affiliated Hospital of China Medical University and the Mental Health Center of Shenyang. Patients with MDD were diagnosed by two trained psychiatrists using the Structured Clinical Interview for DSM-IV Disorders. Participants with MDD met the DSM-IV diagnostic criteria for MDD but not for any other Axis I disorders. The severity of depression was rated using the 17-item HDRS (Williams, 1988) and the Clinical Global Impression of Severity Scale (Guy, 1976). The control group was recruited from the local community. HC participants did not have a current or lifetime history of Axis I disorders or a history of psychotic, mood, or other Axis I disorders in first-degree relatives as determined from the detailed family history. The participants were excluded for the following reasons: 1) a lifetime history of substance/alcohol abuse or dependence, 2) a concomitant major medical disorder, 3) any MRI contraindications, 4) a history of head trauma with loss of consciousness ≥5 minutes or any neurological disorder, and 5) suboptimal imaging data quality. The study was approved by the Institutional Review Board of China Medical University. Among these participants, 1 patient and 5 HCs were excluded because of duplicate data in the data transfer, 1 HC was excluded due to errors in raw DICOM data, 6 patients were excluded due to the use of different scanning parameters, 3 HCs were excluded due to abnormalities in anatomical brain images, 6 patients and 6 HCs were excluded for large head motion during the R-fMRI scan (exceeding 3 mm of translational movements or 3° of rotational movements), 13 patients and 8 HCs were excluded because the MRI scan did not cover the entire brain, 4 patients and 2 HCs were excluded due to a change in the diagnosis in follow-up interviews, 1 HC was excluded due to a lack of demographic information, and 1 HC was excluded because of his young age. Finally, data from the remaining 125 patients with MDD and 249 HCs were used in the present study.

***CSU dataset:*** Patients with MDD were recruited from inpatient or outpatient departments of the Psychiatry Hospital of Zhumadian, Henan Province, China. The diagnosis of MDD was confirmed by two well-trained psychiatrists using the Structured Clinical Interview of the Diagnostic and Statistical Manual of Mental Disorders IV (DSM-IV) criteria for MDD. All HCs were recruited by posting flyers in the local area and had no history of childhood maltreatment. Inclusion criteria for patients with MDD were acutely depressed, medication-free for at least 2 weeks, and depression severity with a minimum score of 20 on the 24-item Hamilton Depression Rating Scale (HDRS) (Williams, 1988). The exclusion criteria for the patients were the presence of a comorbid Axis I disorder, an Axis II personality disorder, and a personal or family history of bipolar disorder. The requirements for HCs were HDRS scores less than 7, no current or history of psychiatric disorders, and no family history of psychiatric disorders. In both groups, additional exclusion criteria included substance abuse or dependence, a neurological or internal illness, and any contraindications to MRI scans. All participants provided both written and verbal informed consent. The study was approved by the Human Investigation Committee of the Second Xiangya Hospital, Central South University of China. Initially, 345 participants, including 217 patients with MDD and 128 HCs, were enrolled. Sixty participants were excluded during quality control, including 8 patients with MDD due to incomplete resting-state functional magnetic resonance imaging (R-fMRI) scans, 1 HC due to errors in raw Digital Imaging and Communications in Medicine (DICOM) data, 10 patients with MDD and 2 HCs due to abnormalities in anatomical brain images, 18 patients with MDD and 17 HCs due to large head motion in R-fMRI scans (exceeding 3 mm of translational movements or 3° of rotational movements), and 4 patients with MDD due to a lack of demographic information. Therefore, data from 177 patients with MDD and 108 HCs from this center were included in the present study.

***PKU dataset:*** Seventy-six drug-naïve patients with first-episode MDD were recruited from the outpatient clinic of the psychiatric departments, and 74 age- and sex-matched HCs were recruited from the local community. The MDD diagnosis was confirmed by two trained psychiatrists using the Mini-International Neuropsychiatric Interview (MINI) (Sheehan et al., 1998), a structured clinical interview for DSM-IV. Inclusion criteria for patients with MDD were a current, acute depressive episode; severe MDD as defined by a minimum score of 24 on the 17-item HDRS (Hamilton, 1967); and a current depressive episode lasting between ≥1 month and ≤24 months. The exclusion criteria for patients with MDD were the presence of a concurrent, comorbid Axis I disorder; an Axis II personality disorder, intellectual disabilities, and any previous or current use of psychotropic medications. The requirements for HCs were no lifetime history of psychiatric disorders, no history of psychiatric disorders in their first-degree relatives, and no history of use of psychotropic medications. All HCs had an HDRS score less than 7, indicating that no HCs suffered from depressive symptoms. An additional questionnaire was used to ensure that all HCs had no experiences that might affect their mood, such as exams, unemployment, and family bereavement, within six months before and during the study. Commonly used exclusion criteria for both the MDD and HC groups were an age of less than 18 or more than 60 years; the presence of an unstable medical condition, neurological illnesses, substance dependence or abuse, or acute suicidal tendencies; and any contraindications to MRI scans. The study was approved by the local Institutional Review Boards. Written informed consent was obtained from all subjects before participation in the present study. The patients voluntarily participated in the study, and no patients who were involuntarily detained in the hospital were included. Data from 1 patient with MDD and 1 HC were excluded because the scans did not cover the entire brain. Therefore, data from 75 patients with MDD and 73 HCs from this center were included in the present study.

***SCU dataset:*** We recruited 52 patients with MDD from the Department of Psychiatry at West China Hospital of Sichuan University. All patients were interviewed using the Structured Clinical Interview for

DSM-IV Disorders by experienced psychiatrists and met the DSM-IV criteria for MDD. The patients were medication-free for at least 2 weeks before scanning and had a 17-item HDRS (Hamilton, 1967) total score of 18 or higher on the day of their MRI scan. The exclusion criteria were as follows: 1) a diagnosis of any psychotic disorders other than MDD; 2) substance or alcohol abuse or dependence within the past 12 months; 3) a history of a significant medical or neurological illness, including any history of significant head trauma with loss of consciousness; and 4) any likelihood that a suicide attempt caused structural abnormalities in the brain. Forty-four HCs were recruited from the local community by a poster advertisement and assessed with the Structured Clinical Interview for the DSM-IV, Non-patient Edition. Exclusion criteria included the same medical and psychiatric factors used to recruit patients, as well as a history of any DSM-IV Axis I disorder or a known history of significant psychiatric illness or suicide among first-degree relatives. All participants were of Chinese Han nationality. The West China Hospital Clinical Trials and Biomedical Ethics Committee of Sichuan University approved our study protocol, and written informed consent was obtained from all participants. Data from 2 patients and 3 HCs were excluded for large head motion during the R-fMRI scan (exceeding 3 mm of translational movements or 3° of rotational movements); thus, 50 patients with MDD and 41 patients from this center were used in the subsequent analyses.

***SWU dataset:*** Patients with MDD were recruited from the outpatient department of the First Affiliated Hospital of Chongqing Medical School in Chongqing, China. All patients were diagnosed by independent assessments of two psychiatrists according to the Structured Clinical Interview for DSM-IV. Disease severity was assessed using the HDRS (Hamilton, 1960) and Beck Depression Inventory (BDI). Before the investigation, we excluded individuals who were not suitable for MRI scanning using an interview and a self-reported checklist. MRI-related exclusion criteria included subjects with claustrophobia, metallic implants, Meniere's syndrome and a history of fainting within the previous 6 months. The exclusion criteria for both groups were current psychiatric disorders (except for MDD) and neurological disorders, substance abuse, and stroke or serious encephalopathy. Notably, all subjects in the control group did not meet the DSM-IV criteria for any psychiatric disorders and did not use any drugs that could affect brain function. This study was approved by the Research Ethics Committee of the Brain Imaging Center of Southwest University and First Affiliated Hospital of Chongqing Medical School. Written informed consent was obtained from each subject. Data from 282 patients with MDD and 254 HCs in this center were included in the subsequent analyses.

Together, after quality control of both clinical information (lack of demographic data, uncertainty in diagnosis and abnormality in brain anatomy) and imaging data (error in Digital Imaging and Communications in Medicine (DICOM) data, incomplete scanning and excessive head motion), the final analysis included 1,434 participants: 709 patients with MDD and 725 HCs.

**Table S1. Clinical information of the subgroups of MDD patients.**

| Subgroup | Duration of illness, mean (SD), yr | Onset age, mean (SD), yr | Episode number, mean (SD) | HDRS, mean (SD) | In first episode (Yes/No) | Medication (Yes/No) |
|---|---|---|---|---|---|---|
| First episode (n=412) | 1.86 (3.23) (n=387) | 29.35 (10.18) (n=181) | 1 (0) (n=412) | 22.13 (6.40) (n=402) | 412/0 | 131/274 |
| Non-first episode (n=80) | 7.63 (7.01) (n=74) | 25.00 (6.86) (n=9) | 2.50 (.69) (n=80) | 20.66 (5.74) (n=79) | 0/80 | 60/18 |
| Statistics T or $\chi^2$/P | 11.18/<.001 | 1.27/.21 | 22.51/<.001 | 1.90/.06 | N.A. | 54.36/<.001 |
| With medication (n=198) | 4.25 (5.49) (n=191) | 27.16 (9.69) (n=46) | 2.25 (.85) (n=24) | 19.94 (6.63) (n=192) | 131/60 | 190/0 |
| Without medication (n=301) | 1.71 (3.47) (n=273) | 30.05 (10.11) (n=151) | 1 (0) (n=99) | 23.32 (5.95) (n=295) | 274/18 | 0/301 |
| Statistics T or $\chi^2$/P | 6.11/<.001 | 1.71/.09 | 14.88/<.001 | 5.85/<.001 | 54.36/<.001 | N.A. |
| Onset age >21 (n=221) | 1.56 (2.76) (n=221) | 33.39 (7.75) (n=221) | 1 (0) (n=93) | 23.09 (6.97) (n=149) | 138/6 | 33/117 |
| Onset age ≤21 (n=63) | 2.38 (4.37) (n=63) | 17.47 (2.72) (n=63) | 1 (0) (n=10) | 20.89 (8.18) (n=44) | 43/3 | 13/34 |
| Statistics T or $\chi^2$/P | 1.81/.07 | 16.01/<.001 | N.A. | 1.77/.08 | .43/.51 | .64/.42 |

Abbreviations: SD, standard deviation; HDRS, Hamilton depression rating scale; N.A., not available.

**Figure S1. Effect of age on functional measurements.** The figure illustrates significant effects of age on functional measurements from local voxels to distant brain connections using a stepwise regression analysis. Warm and cold colors indicate positive and negative correlations between functional coordination and age, respectively. The surface rendering was created using BrainNet Viewer (www.nitrc.org/projects/bnv/) (Xia et al., 2013). ALFF, amplitude of low-frequency fluctuations; ReHo, regional homogeneity; FCS, functional connectivity strength; GRF, Gaussian random field.

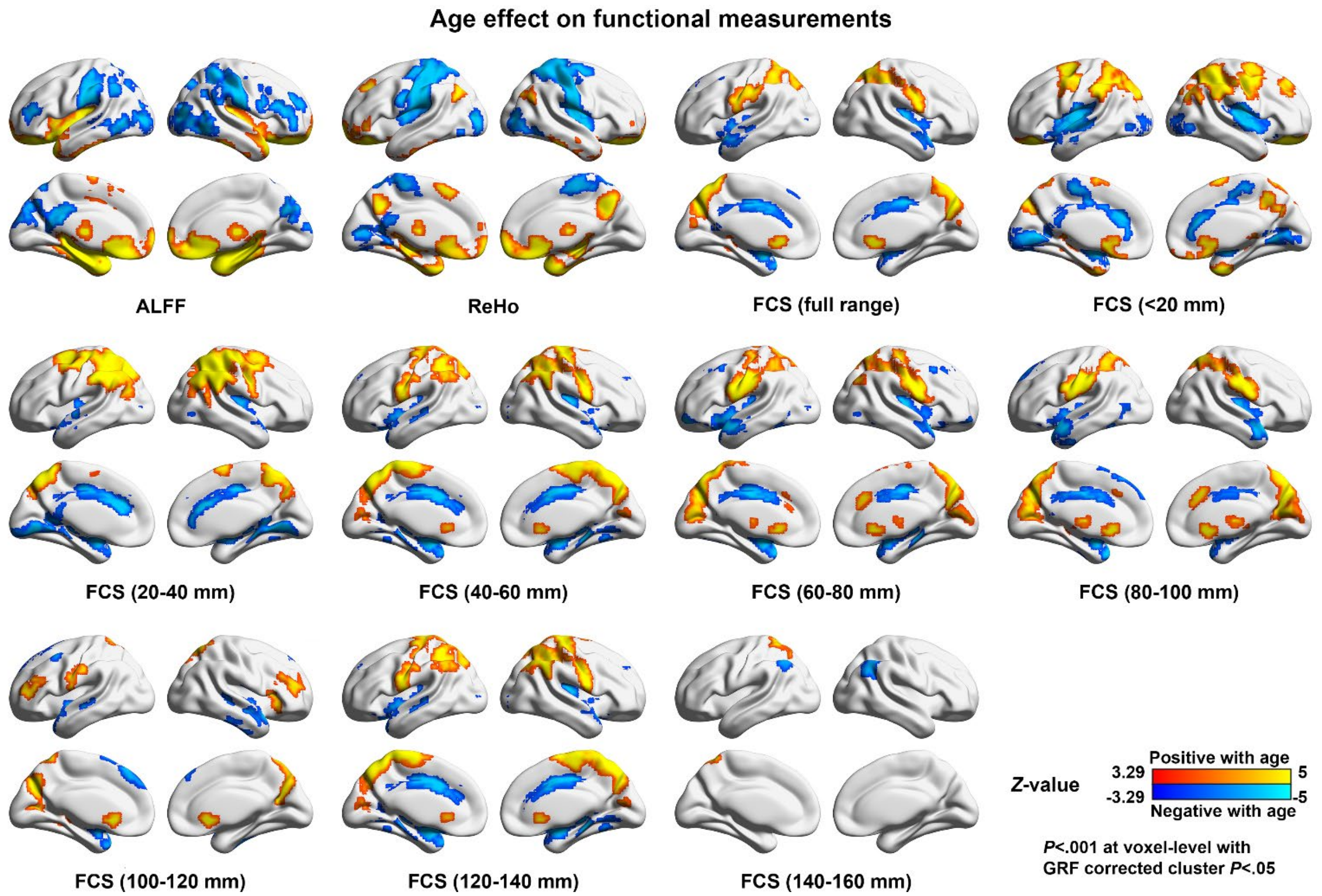

**Figure S2. Effect of sex on functional measurements.** The figure illustrates significant effects of sex on functional measurements from local voxels to distant brain connections using a stepwise regression analysis. Warm and cold colors indicate positive and negative correlations between functional coordination and age, respectively. ALFF, amplitude of low-frequency fluctuations; ReHo, regional homogeneity; FCS, functional connectivity strength; GRF, Gaussian random field.

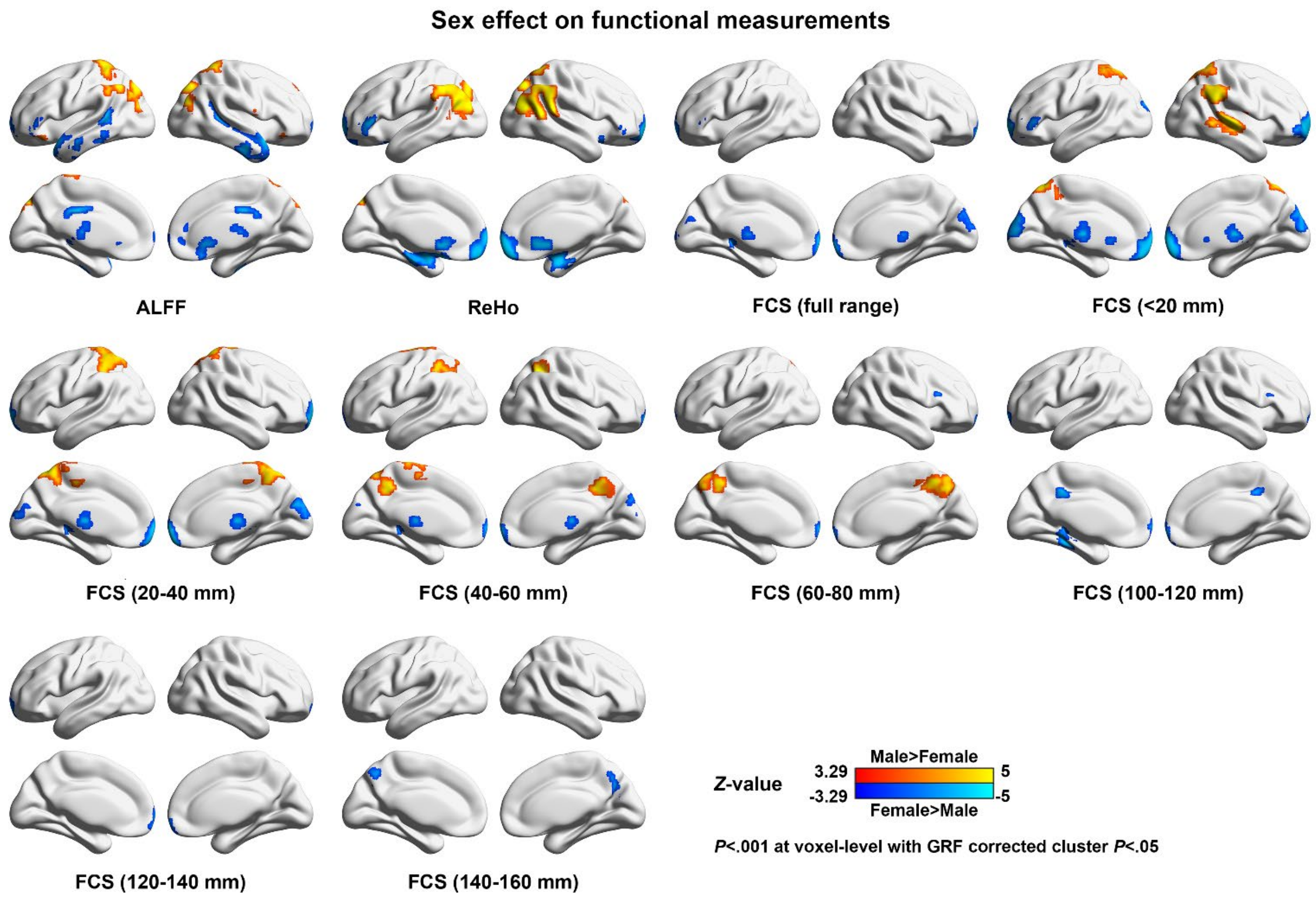

**Figure S3. Differences in functional measurements between patients with MDD and healthy controls using data filtered with a band of 0.01-0.08 Hz.** The figure illustrates significant between-group effects on functional measurements from local voxel to distant brain connections using the stepwise regression analysis and Liptak-Stouffer *z*-score method on data with and without GSR. Warm and cold colors indicate higher and lower functional measurements in patients with MDD than in the HCs, respectively. MDD, major depressive disorder; HC, healthy controls; s-w, stepwise; nGSR, nonglobal signal regression; L-S, Liptak-Stouffer; GSR, global signal regression; ALFF, amplitude of low-frequency fluctuations; ReHo, regional homogeneity; FCS, functional connectivity strength; GRF, Gaussian random field.

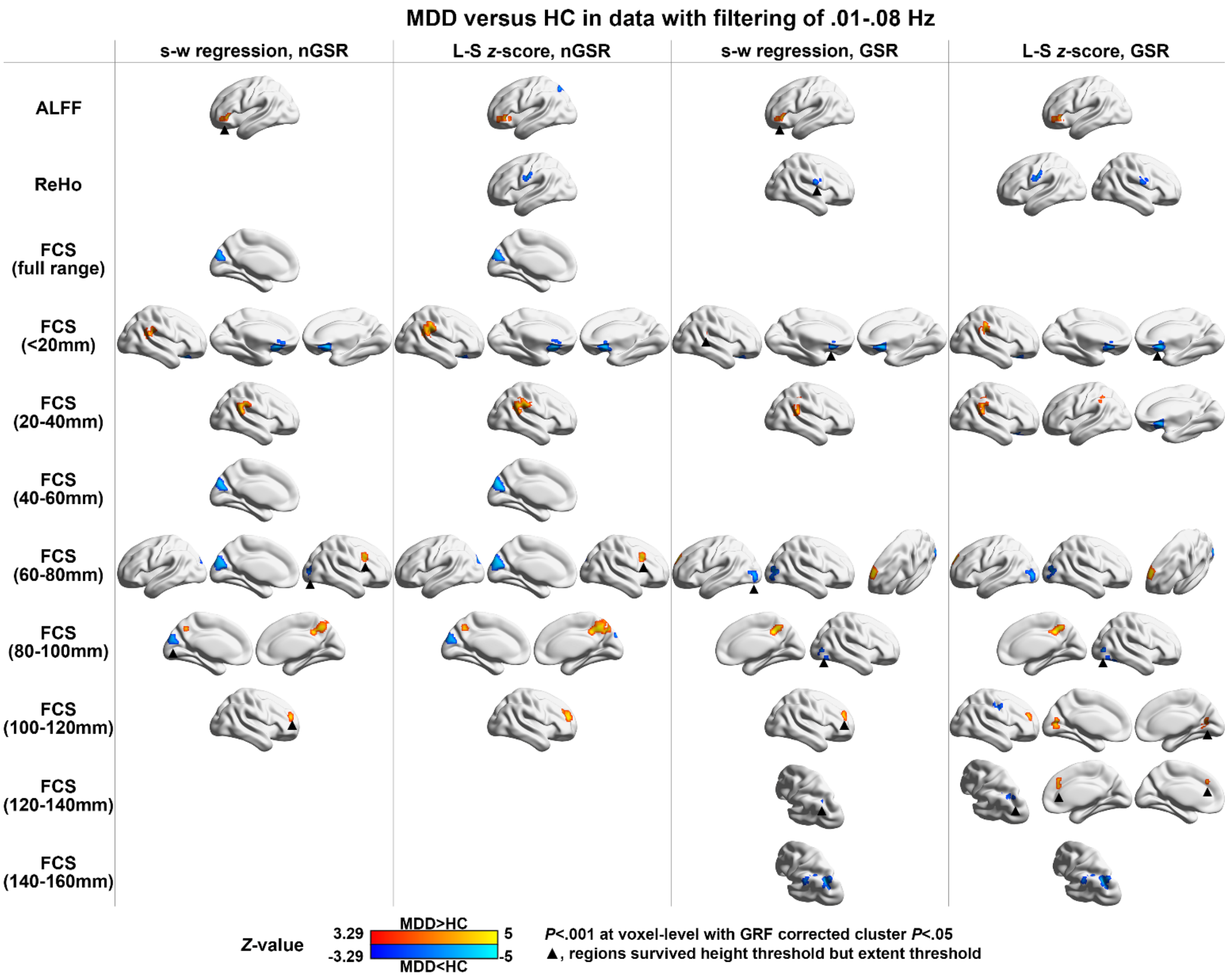

**Figure S4. Site effect on functional measurements.** The figure illustrates significant site effects on functional measurements from local voxel to distant brain connections using the Kruskal-Wallis tests. ALFF, amplitude of low-frequency fluctuations; ReHo, regional homogeneity; FCS, functional connectivity strength; GRF, Gaussian random field.

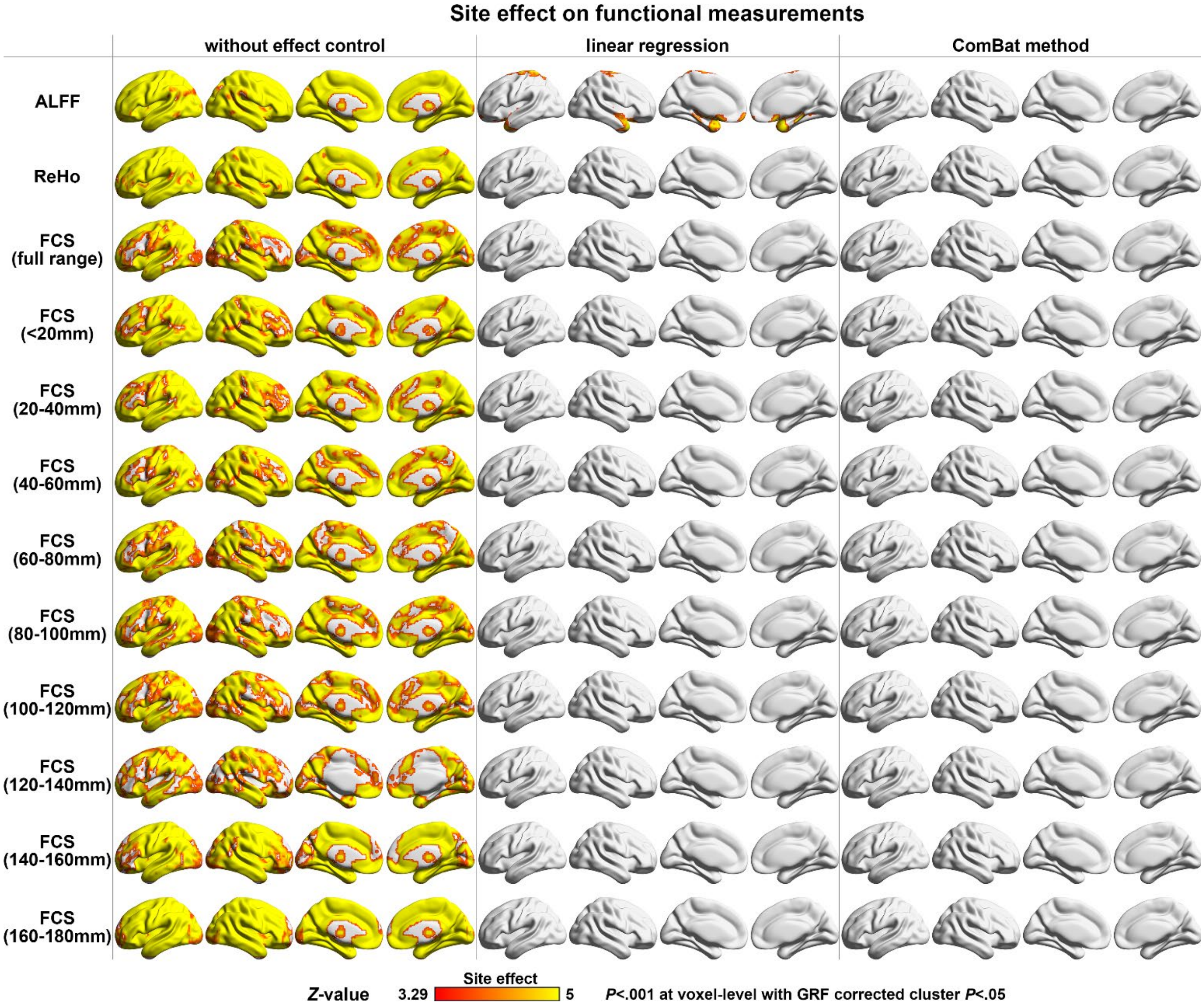

**Figure S5. Differences in functional measurements between patients with MDD and healthy controls using ComBat to reduce the site effect.** The figure illustrates significant between-group effects on functional measurements from local voxel to distant brain connections using the stepwise regression analysis on data with and without GSR. Warm and cold colors indicate higher and lower functional measurements in patients with MDD than in the HCs, respectively. MDD, major depressive disorder; HC, healthy controls; s-w, stepwise; nGSR, nonglobal signal regression; GSR, global signal regression; ALFF, amplitude of low-frequency fluctuations; ReHo, regional homogeneity; FCS, functional connectivity strength; GRF, Gaussian random field.

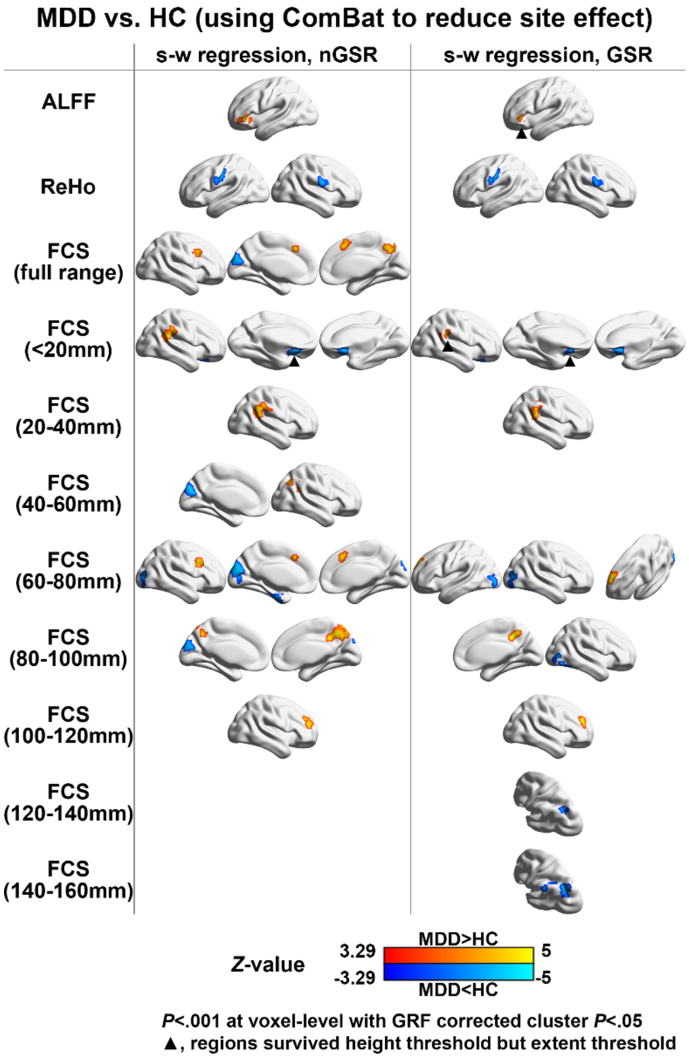

**Figure S6. Robust fitting of functional measurements and clinical variables in each center.** Each dot represents a subject, and its color indicates its weight in the robust regression analysis. A color map of blue to gray indicates the regression weight from high to low, respectively. Dashed lines indicate the confidence interval of the regression. The functional measurements and clinical variables were fitted by age, sex and centers. CSU, Central South University; SCU, Sichuan University; CMU, China Medical University; R.IFGOperc, right inferior frontal gyrus, opercular part; R.SMG, right supramarginal gyrus; R.PoCG, postcentral gyrus; FCS, functional connectivity strength; HDRS, Hamilton depression rating scale.

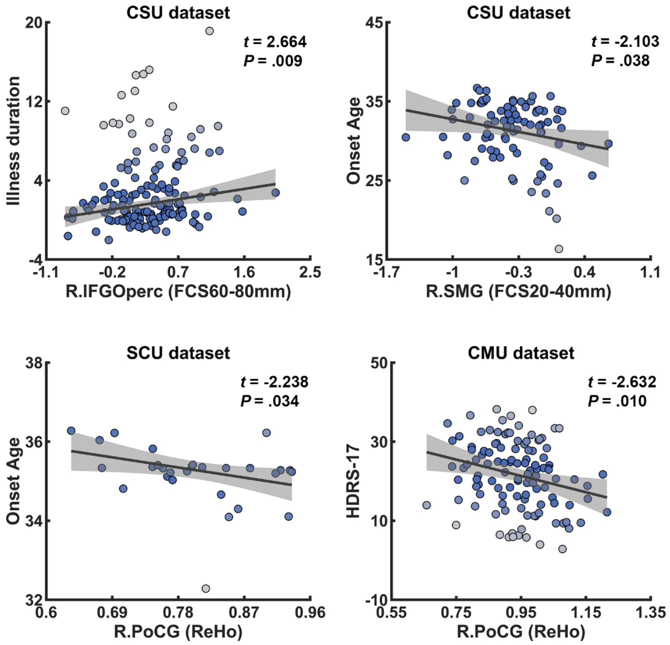
